# N4BP1 is dimerization-dependent linear ubiquitin reader regulating TNFR1 signalling through linear ubiquitin binding and Caspase-8-mediated processing

**DOI:** 10.1101/2021.11.02.466974

**Authors:** Katarzyna Kliza, Wei Song, Irene Pinzuti, Simone Schaubeck, Simone Kunzelmann, David Kuntin, Arianna Fornili, Alessandro Pandini, Kay Hofmann, James A. Garnett, Benjamin Stieglitz, Koraljka Husnjak

## Abstract

Signalling through TNFR1 modulates proinflammatory gene transcription and programmed cell death, and its impairment causes autoimmune diseases and cancer. NEDD4-binding protein 1 (N4BP1) was recently identified as a critical suppressor of proinflammatory cytokine production^1^, whose mode of action remained unknown. Here, we show that N4BP1 is a novel linear ubiquitin reader that negatively regulates NFκB signalling by its unique dimerization-dependent ubiquitin-binding module that we named LUBIN. Dimeric N4BP1 strategically positions two non-selective ubiquitin-binding domains to ensure exclusive recognition of linear ubiquitin. Under proinflammatory conditions, N4BP1 is recruited to the nascent TNFR1 signalling complex, where it regulates duration of proinflammatory signalling in LUBIN-dependent manner. N4BP1 deficiency accelerates TNFα-induced cell death by increasing complex II assembly. Under proapoptotic conditions, Caspase-8 mediates proteolytic processing of N4BP1 and the resulting cleavage fragment of N4BP1, which retains the ability to bind linear ubiquitin, is rapidly degraded by the 26S proteasome, accelerating apoptosis. In summary, our findings demonstrate that N4BP1 dimerization creates a unique linear ubiquitin reader that ensures timely and coordinated regulation of TNFR1-mediated inflammation and cell death.

## Main text

TNFα is a potent, pleiotropic cytokine, regulating inflammation, immunity and programmed cell death^2^. Activation of TNFR1 induces the assembly of the membrane-bound TNFR1 signalling complex (TNFR1-SC) composed of adaptor proteins (i.e., TRADD), kinase RIP1 and E3 ligases, including LUBAC^3^. TNFR1 signalling highly depends on numerous posttranslational modifications, including linear ubiquitination^3,4^. The E3 ligase complex LUBAC stabilizes TNFR1-SC by assembling linear ubiquitin (Ub) chains on several complex components (TNFR1, RIP1, NEMO). These modifications are recognized by a subset of linear Ub-binding domain (UBD)-containing (LUBID) proteins that further regulate downstream signalling, including the UBAN (Ub binding in ABIN and NEMO) domain in NEMO^4–8^. Together with K11-, K48- and K63-linked Ub chains^7,9^, these Ub linkages provide a platform for the recruitment of the kinase complex IKK to initiate NFκB pathway for pro-survival gene induction^3,10^. Prolonged activation of TNFR1 signalling can also trigger the assembly of the cytosolic complex II, composed of TRADD, FADD, RIP1, c-FLIP and procaspase-8, subsequently leading to caspase-8 (CASP8) activation^11^. Among others, apoptosis progression requires removal of M1 linkages from FADD and linear Ub-modified substrates within TNFR1-SC^4,12^. Moreover, LUBAC deficiencies promote complex II assembly and induce aberrant TNFα-mediated endothelial cell death^13^.

## N4BP1 is a novel linear ubiquitin reader

By using a Y2H assay, we identified a novel UBD in N4BP1 and demonstrated that endogenous N4BP1 selectively binds linear Ub **(Fig. 1A).** The minimal UBD obtained by Y2H encompasses the divergent CUE domain of N4BP1 **(Fig. 1B)**. Bioinformatic analysis identified two additional putative UBDs in N4BP1: non-functional UBM-like domain and divergent UBA domain **(Fig. 1B, Extended Data Figs. 1A-1B)**, which bind to all Ub species **(Extended Data Fig. 1A)**. However, our Ub-binding analysis indicated that N4BP1’s divergent CUE domain is indispensable for specific recognition of linear Ub chains **(Extended Data Fig. 1A)**. Accordingly, mutation of the canonical Ub-binding motif, FP (F862G/P863A in mouse N4BP1), which is conserved among CUE domains **(Extended Data Fig. 1C)**^14–16^, abolished the binding between the isolated CUE domain and linear tetraUb **(Fig. 1C**). In order to further investigate the Ub-binding specificity of N4BP1 for linear Ub, we performed ITC measurements and determined K_D_ values for the isolated CUE domain **(Table 1, Extended Data Fig. 2A–2E**). Since the obtained equilibrium dissociation constants did not indicate a clear preferential interaction between N4BP1 and M1-linked Ub chains, we characterised the binding by NMR spectroscopy and established a structural model for the N4BP1 CUE domain spanning residues 850-893. The obtained structure folds into the canonical three-helical bundle, as observed for other CUE domains^17^.

**Figure 1.**
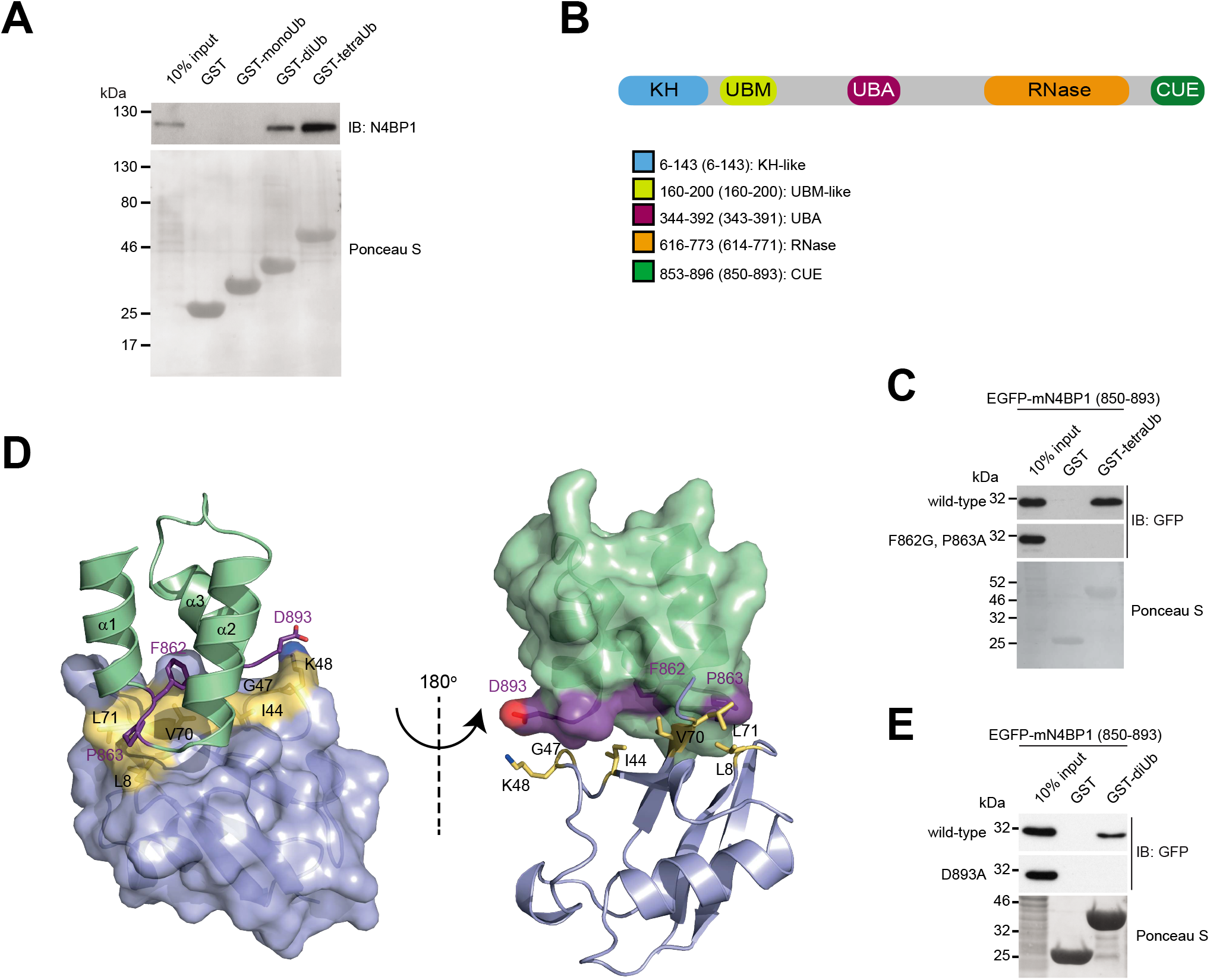
N4BP1 is a novel linear Ub-binding protein. (**A**) GST pull-down assay of endogenous mouse N4BP1 with GST fusions of mono-, di- and tetraUb. (**B**) Schematic representation of human N4BP1 with predicted domains. Domains: KH-like (K Homology), UBM-like (Ub-binding motif-like), divergent UBA (Ub-associated), NYN RNase and divergent CUE. Legend: the numbers outside and inside the brackets indicate amino acid residues of human and mouse N4BP1 respectively. (**C**) GST pull-down assay with GST fusions of mono- and tetraUb with N4BP1 CUE domain and predicted Ub binding-deficient (F862G, P863A) mutant transiently overexpressed in HEK293T cells as EGFP fusions. (**D**) Structural model of the CUE domain of N4BP1 (green) in the complex with Ub (blue) in a cartoon and surface presentation. Residues of N4BP1 and Ub, which form the interface of the complex are shown in purple and yellow respectively. (**E**) GST pull-down assay of mouse N4BP1 CUE domain (850-893) transiently overexpressed as EGFP fusion in HEK293T cells with GST fusions of mono- and diUb. The mutation D893A diminishes Ub binding by disrupting the polar interaction between N4BP1 and Ub, as indicated in the structural model of the complex in **D**.

**Table 1.** K_D_ values of different N4BP1 fragments with various ubiquitin linkages determined by ITC and SPR measurements.

| ITC | | $K_D$<br>[μM] | $\Delta H$<br>[kcal/mol] | $-T\Delta S$<br>[kcal/mol] | $\Delta G$<br>[kcal/mol] | N |
| --- | --- | --- | --- | --- | --- | --- |
| N4BP1 (850-893) | monoUb | $27.5 \pm 5.6$ | -5.2 | -1.03 | -6.21 | 0.915 |
| | M1-diUb | $28.1 \pm 2.7$ | -5.3 | -0.86 | -6.21 | 2.16 |
| | K63-diUb | $16.7 \pm 8.2$ | -5.88 | -3.15 | -6.62 | 1.71 |
| | K48-diUb | $47.0 \pm 5.6$ | -5.87 | -0.42 | -5.67 | 1.29 |
|  | monoUb K48A | no binding detected | - | - | - | - |
| | monoUb K48R | $28.1 \pm 9$ | -4.61 | -1.58 | -6.2 | 0.82 |

| SPR | | $K_D$<br>[μM] |
| --- | --- | --- |
| N4BP1 (613-774) | monoUb | no binding detected |
|  | M1-diUb |  |
|  | K63-diUb |  |
|  | K48-diUb |  |
| N4BP1 (613-893) | monoUb | 84.93 |
|  | M1-diUb | 0.43 |
|  | K63-diUb | 10.85 |
|  | K48-diUb | 34.23 |

To reveal how the CUE domain forms distinctive interfaces with different Ub linkages, we performed ^1^H-^15^N HSQC NMR titration experiments with monoUb, K48-, K63- and M1-linked diUb (**Extended Data Figs. 3A–3B, 4A–4B**). All interface areas between the CUE domain and Ub species are highly similar, confirming that the isolated CUE domain of N4BP1 recognizes Ub in a nonspecific manner. To further investigate how the CUE domain interacts with Ub, we found that it forms a small hydrophobic interface of 830 Å^2^ with Ub, which comprises the aliphatic portion of the C-terminal D893 of helix α3 and F862 and P863 residues located between helices α1 and α2 (**Fig. 1D**) and recognizes a nonpolar surface area of Ub, involving the hydrophobic patch around I44. Chemical shift mapping of monoUb, K63- or M1-linked diUb show marked chemical shift perturbation (CSP) values for K48 residue of Ub, which is absent in K48-linked diUb (**Extended Data Fig. 4C**) and contributes to the interface by forming a contact with D893 of the CUE domain of N4BP1. Mutations D893A in N4BP1 or K48A in Ub, abolish complex formation, indicating that this polar interaction is essential for robust binding (**Fig. 1E**, **Table 1**, **Extended Data Fig. 2**). Interestingly, mutation K48R in Ub does not affect the affinity towards N4BP1, which demonstrates that a positive charge at the position K48 plays a dominant role in the interaction with N4BP1, a finding that is in line with the reduced ability of the isolated CUE domain to co-precipitate K48-diUb (**Extended Data Figs. 5A–5B**). Here, the side chain of the K48 participates in the formation of the isopeptide bond, and therefore, only the distal Ub moiety of K48-linked Ub chains can fully engage with the CUE domain of N4BP1, causing a reduced binding affinity compared to other Ub linkages.

## An N4BP1 dimer functions as a unique linear ubiquitin reader

We set out to determine how specificity for Met1-linked Ub chains is achieved. Towards that aim, we tested three CUE-containing N4BP1 fragments (aa850-893, aa706-893 and aa343-893) for binding to a set of synthetic diUbs^18–20^ and observed increasing specificity for the linear diUb (**Extended Data Figs. 5A–5B**). We therefore hypothesized that the adjacent RNase domain is involved in the high affinity binding to M1-linked Ub chains and probed this interaction by SPR measurements. Remarkably, the extended construct that comprised the RNase and CUE domains (aa613-893) shows a 65-fold increase in linear Ub binding, while all other tested Ub interactions display no significant difference when compared to the isolated CUE domain (**Fig. 2A**, **Table 1**). However, we did not observe any interaction between the isolated RNase domain (613-773) and Ub linkages (**Table 1, Extended Data Fig. 5C**), further raising the question of how N4BP1 specificity for Met1-linked Ub chains is achieved. Intriguely, N4BP1 constructs comprising the RNase domain had a strong tendency to self-associate *in vivo* and *in vitro* (**Extended Data Figs. 6A–6E**), suggesting that N4BP1 dimerizes through its RNAse domain. Interestingly, the protein MCPIP1, which shares 52% sequence identity with N4BP1, features the same oligomeric nature caused by the formation of an asymmetric dimer of its RNase domain^21^. A homology model of the N4BP1 RNase domain, based on the structure of the MCPIP1 RNase domain, shows that an N4BP1 dimer can be formed (**Extended Data Fig. 6F**). We confirmed the N4BP1 dimerization mode by creating point mutants of several interface residues and found that these mutations completely abolished N4BP1 dimerization (**Fig. 2B**). To explain how N4BP1 dimerization creates specificity for linear linkages, we hypothesized that the spatial arrangement of the dimer brings the two adjacent CUE domains into close vicinity, which could result in the simultaneous interaction of the CUE domains with the same linear M1-linked diUb molecule (**Fig. 2C, Extended Data Fig. 6G**). To further examine this scenario, we deployed a homology modelling and docking approach to generate a model of N4BP1 (613-893) in complex with M1-linked diUb, which aligns with the experimental constraints derived from our CSP experiments and mutational analysis of interface residues between the CUE domain and Ub. The refined model, which satisfies all experimental criteria, demonstrates that dimerization of N4BP1 *via* the RNase domain is well suited to generate a spatial composite arrangement of the CUE domains that permits binding of M1-diUb with high affinity (**Fig. 2D)**

**Figure 2.**
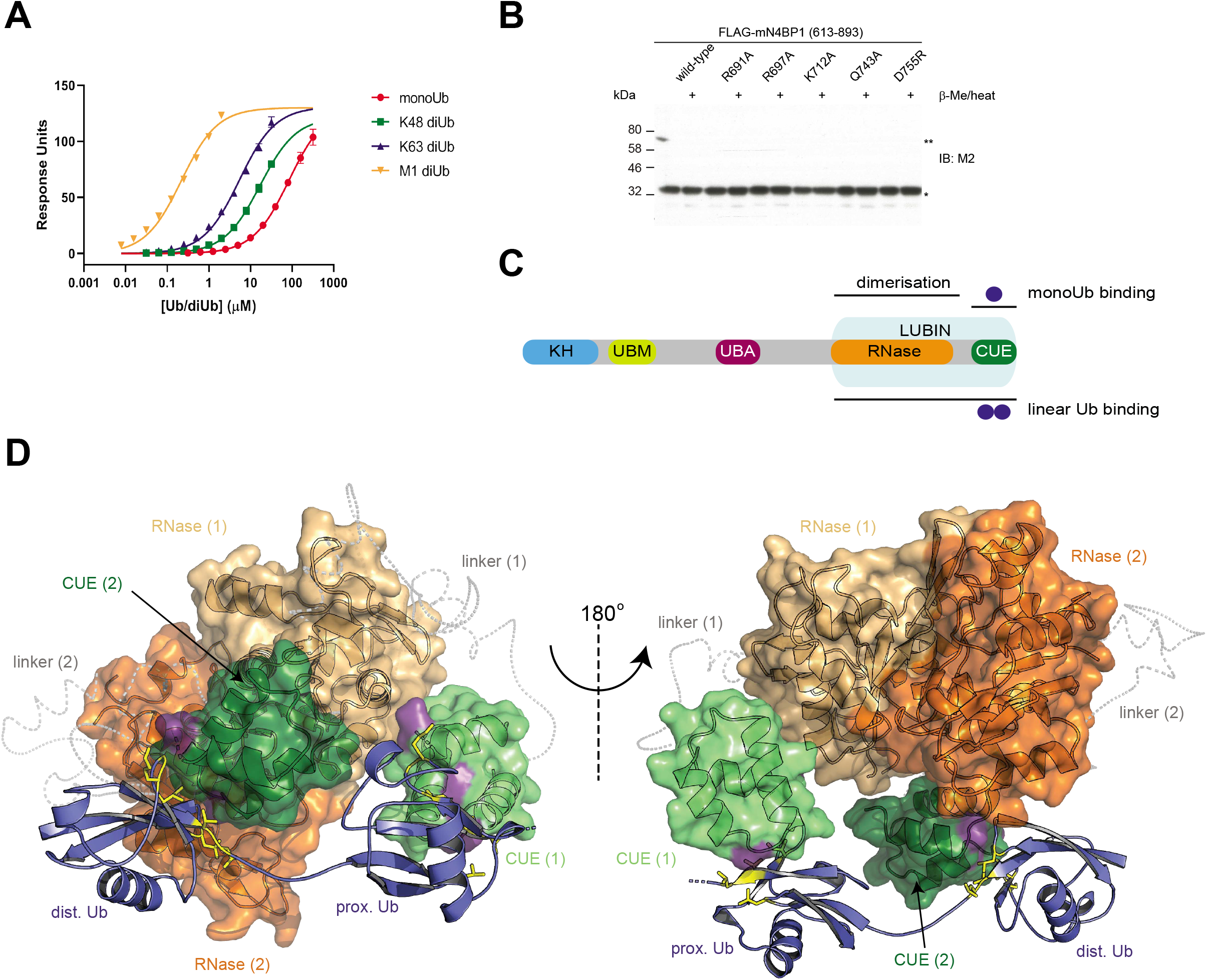
N4BP1 achieves linear ubiquitin binding through unique dimerization of its RNase and CUE domains. (**A**) Quantitative analysis of Ub binding specificity of N4BP1 (613-893) by surface plasmon resonance (SPR). Average binding responses of increasing concentrations of monoUb and K48-, K63- and M1-linked diUb were fitted to a saturation equilibrium binding model to obtain equilibrium dissociation constants (**B**) The effect of point mutations in the RNase domain on dimerization of mouse N4BP1 (613-893). Each lysate was prepared both under non-denaturing (lysate mixed with 1xLDS buffer) or denaturing (lysate mixed with 1xLDS+ β-Me, followed by thermal denaturation) conditions and analysed by Western blotting. (*) indicates N4BP1 (613-893) monomer, (**) is N4BP1 (613-893) dimer that is destroyed under denaturing conditions. (**C**) Schematic representation of human N4BP1 with indicated monoUb- and linear Ub-binding domains, as well as dimerization module consisting of RNase and CUE domains. (**D**) Structural model of the N4BP1 (613-893) dimer in complex with M1-linked diUb. A homology model of the RNase domain was generated using residues 135-339 of the X-ray structure of dimeric MCPIP1 (PDB ID: 5H9W) as a template. The RNase domain (orange) is interconnected to the NMR based solution structure of the CUE domain (green) *via* a flexible linker region (residues 775-849), which is shown as a C-α backbone trace (grey, dashed line). The dimeric arrangement of N4BP1 is compatible with simultaneous interaction of both CUE domains with the experimentally defined interface areas of M1-linked diUb (blue). Residues involved in N4BP1 recognition are shown in yellow and the corresponding contact surface of the CUE domain is depicted in purple.

To validate the dimer-induced mechanism for high affinity interaction with linear Ub in more detail, we probed the ability of the artificially dissociated RNase-like domain (**Extended Data Fig. 6H**) and RNase-like domain interface mutants to specifically bind to M1-linked diUb (**Extended Data Fig. 6I, Extended discussion**). Strikingly, the capacity of all the mutants to specifically recognize linear Ub was drastically reduced, which further supports the notion that both CUE-like domains of the N4BP1 dimer engage in specific interaction with a single M1-linked Ub chain. Furthermore, a prerequisite to allow simultaneous interaction of both CUE-like domains with linear Ub is the linker region (775-849) that connects RNase- and CUE-like domains and its deletion interferes with the ability of N4BP1 to bind Ub in a linear Ub-specific manner (**Extended Data Fig. 6I**). Since we could not detect a direct intramolecular interaction between the RNase and the CUE domain, which would fix the relative orientation beween both domains (**Extended Data Fig. 2F**), we suggest that the linker region acts as a scafold that supports the specific arrangement of CUE domains in the N4BP1 dimer to establish linear Ub specificity.

In conclusion, our data suggest that N4BP1 is utilising a novel, unique mechanism for M1-linked Ub chains recognition. We therefore named the newly discovered N4BP1 LUBID “Linear Ub-Interacting Domain in N4BP1” (LUBIN) (**Figure 2C**).

## N4BP1 is a negative regulator of NFκB signalling

Basal cellular levels of linear Ub species are very low and their assembly is induced by stimuli, such as TNFα, IL-1β and poly (I:C)^6^. Identification of TNFα-induced linear Ub high molecular weight (HMW) species in immunoprecipitated endogenous N4BP1 samples, further confirmed the ability of N4BP1 to bind linear Ub chains (**Fig. 3A**), and prompted us to examine the role of N4BP1 in TNFα-induced NFκB signalling. Towards that aim, we utilized the N4BP1 knock-out (N4BP1^-/-^) and wild-type (N4BP1^+/+^) mouse embryonic fibroblasts (MEFs) that express similar levels of various proteins involved in TNFR1 signalling pathway (**Extended Data Fig. 7A**). We observed a strong increase of TNFα-induced NFκB transcription activity (**Extended Data Fig. 7B**), the expression of TNFα target genes CXCL1 and IL-6 (**Extended Data Fig. 7C**), and significantly increased nuclear translocation of NFκB subunit p65 (**Fig. 3B, Extended Data Fig. 7D**) in N4BP1^-/-^ MEFs, contrary to N4BP1^+/+^ MEFs. Consistently, kinetics of the IκBα phosphorylation and degradation differ in N4BP1^-/-^ and N4BP1^+/+^ MEFs (**Fig. 3C**, **Extended Data Fig. 7E**). Noteworthy, cytoplasmic N4BP1 (**Extended Data Fig. 7F–7G**)^22^ specifically affects TNFα-induced NFκB pathway, as IκBα degradation kinetics induced by IL-1β stimulation remains unaltered in N4BP1^-/-^, compared to N4BP1^+/+^ MEFs (**Extended Data Fig. 7H**). N4BP1 functions in close proximity to TNFR1, as it is recruited to the nascent TNFR1-SC within 5 min of TNFα stimulation (**Fig. 3D**). To determine the correlation between N4BP1 ability to bind linear Ub chains and its function in proinflammatory TNFR1 signalling, we monitored the activation of NFκB pathway upon TNFα stimulation in N4BP1^-/-^ MEFs reconstituted with either empty vector, HA-N4BP1 (1-893) or HA-N4BP1 (1-893, F862G/P863A) (**Extended Data Fig. 8A**). Contrary to N4BP1^+/+^, Ub binding-deficient N4BP1 mutant failed to restrict activation of NFκB pathway (**Fig. 3E, Extended Data Fig. 8B**). The presence of N4BP1 stabilized TNFα-induced linear Ub HMW conjugates over time, implying the competition between N4BP1 and linear Ub-specific deubiquitinating enzymes (DUBs) for M1 linkages (**Extended Data Fig. 8C**). Hence, N4BP1 is a previously uncharacterized component of TNFR1-SC that modulates proinflammatory TNFR1 signalling through its ability to specifically recognize linear Ub chains.

**Figure 3.**
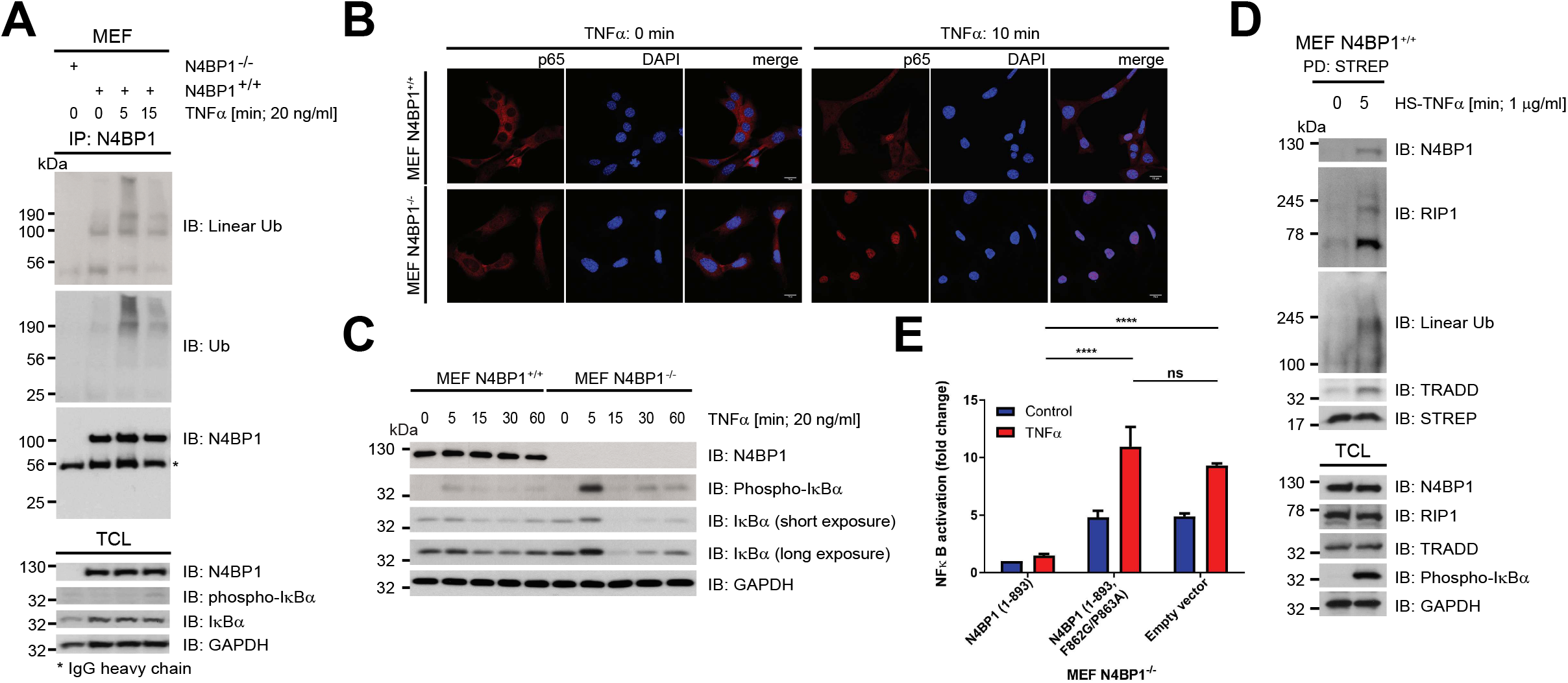
N4BP1 is a novel negative regulator of TNFR1 signalling. (**A**) Binding of M1-linked HMW Ub species to N4BP1. Immunoprecipitated endogenous N4BP1 and total cell lysates upon TNFα (20 ng/ml) treatment for indicated time periods were analysed by Western blotting with indicated antibodies. (**B**) Effect of N4BP1 on nuclear translocation of p65 upon TNFα stimulation. Sixteen hours post-starvation, N4BP1^+/+^ and N4BP1^-/-^ MEFs were treated with TNFα (20 ng/ml) for 10 min. Localization of p65 was detected by indirect immunofluorescence using anti-p65 antibody (1^st^ and 4^th^ column). Nucleoli were stained with DAPI (2^nd^ and 5^th^ column). (**C**) The effect of N4BP1 on activation of TNFα-mediated NFκB pathway. After 16 hours of serum starvation, N4BP1^+/+^ and N4BP1^-/-^ MEFs were treated with TNFα (20 ng/ml) for indicated time periods. Total cell lysates were analysed by Western blotting with indicated antibodies. (**D**) Analysis of N4BP1 interaction with TNFR1-SC complex. After 16 hours starvation, N4BP1^+/+^ MEFs were stimulated with recombinant HS-TNFα (1 μg/ml, 5 min) and TNFα-bound signalling complex was pulled down with Strep-tactin resins and subsequently resolved and analysed by Western blotting with indicated antibodies. (**E**) The effect of linear Ub binding-deficient N4BP1 on NFκB transcriptional activity. N4BP1^-/-^ MEFs stably expressing either empty vector, HA-N4BP1 (1-893) or HA-N4BP1 (1-893, F862G/P863A) were transiently transfected with pNFκB-Luc and pUT651 plasmids encoding luciferase and ß-galactosidase, respectively. After 24 hours, cells were starved for 16 hours, followed by 6 hours stimulation with TNFα (20 ng/ml). Lysates were subjected to luciferase and ß-galactosidase assays. Three independent experimental replicates consisting of technical duplicates were performed. Results are shown as means and s.e.m. (n=3). n.s., no statistically significant difference, * *P* < 0.05 and **** *P* < 0.0001, determined by two-way ANOVA test *post hoc* Sidak’s multiple comparisons test.

## N4BP1 is cleaved by CASP8 upon prolonged TNF stimulation

The presence of M1 linkages within TNFR1-SC preserves the architecture of the complex and prevents assembly of the complex II, which is a prerequisite for apoptotic cell death^3^. As such, we next examined the effect of N4BP1 on TNFα-mediated cell death. N4BP1-deficient MEFs were significantly sensitized to apoptosis induced by combined TNFα and CHX treatment (**Fig. 4A–4B**). The suppressive effect of N4BP1 on apoptosis depends on a functional LUBIN, as N4BP1^-/-^ MEFs reconstituted with HA-N4BP1 (1-893, F862G/P863A) are significantly more susceptible to cell death than N4BP1^-/-^ MEFs stably expressing HA-N4BP1 (1-893) (**Fig. 4C**). Accordingly, N4BP1 deficiency enhanced the assembly of RIP1 and proximal components of the complex II (FADD and CASP8) under proapoptotic conditions (**Fig. 4D**, lanes 5 and 6). Moreover, N4BP1 was found in complex II upon cell death-inducing treatments (**Fig. 4D**, lanes 3 and 5). FADD is modified by LUBAC and deubiquitinated upon apoptosis induction^12^. Hence, the observed recruitment of N4BP1 to complex II could be explained by the recognition of linear Ub-modified FADD by N4BP1 LUBIN, which presumably sequesters FADD in TNFR1-SC, thereby slowing down the complex II formation.

**Figure 4.**
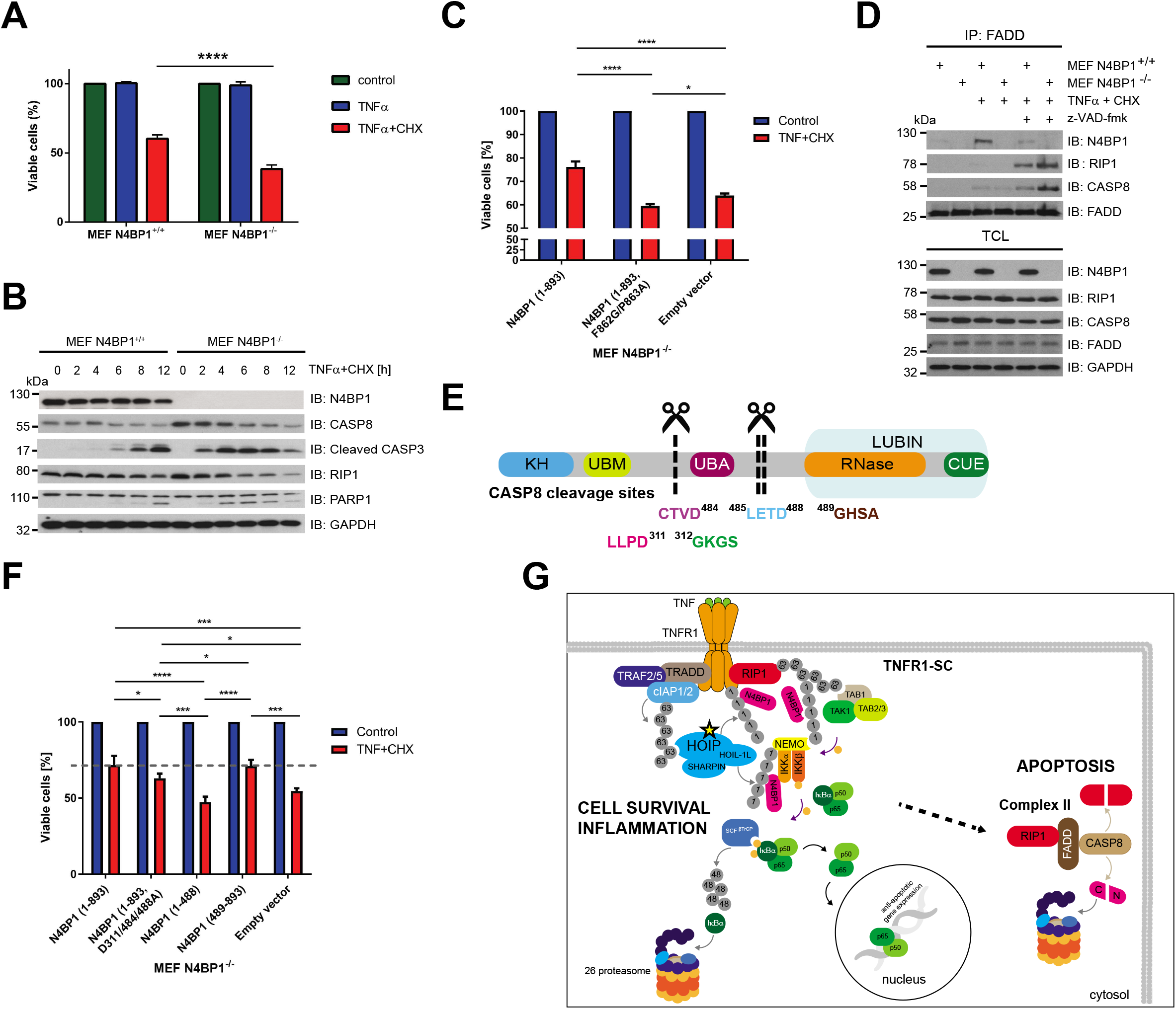
CASP8-mediated proteolytic processing of N4BP1 promotes TNFα-induced cell death. (**A**) Cell viability of N4BP1^+/+^ and N4BP1^-/-^ MEFs left untreated or exposed for 12 hours to TNFα alone (10 ng/ml) or TNFα combined with CHX (10 ng/ml and 0.5 μg/ml respectively). Cell survival was determined by crystal violet staining. Results are shown as means and s.e.m. (n=3). **** *P* < 0.0001, determined by two-way ANOVA, *post hoc* Sidak’s multiple comparisons test. (**B**) The effect of N4BP1 on apoptotic cell death. N4BP1^+/+^ and N4BP1^-/-^ MEFs were treated with TNFα (10 ng/ml) and CHX (0.5 μg/ml) for indicated time periods. The proteolytic processing of apoptotic markers was measured by monitoring the appearance of a cleaved form of CASP3, the disappearance of the full-length CASP8 and RIP1, and both cleaved and uncleaved forms of PARP1. (**C**) Cell viability of N4BP1^-/-^ MEFs reconstituted with empty vector, N4BP1 (1-893) or N4BP1 (1-893, F862G/P863A) under apoptotic conditions. Cells were left untreated or exposed for 10 hours to TNFα and CHX (10 ng/ml and 0.5 μg/ml respectively). Cell viability was determined by the CellTiter-Glo Luminescent assay. Three independent experimental replicates consisting of technical triplicates were performed. Results are shown as means and s.e.m. (n=3). n.s., no statistically significant difference, * *P* < 0.05, **** *P* < 0.0001, determined by two-way ANOVA, *post hoc* Sidak’s multiple comparisons test. (**D**) Effect of N4BP1 on proapoptotic complex II formation. N4BP1^+/+^ and N4BP1^-/-^ MEFs were left untreated or exposed to TNFα and CHX (10 ng/ml and 0.5 μg/ml, respectively) without or with z-VAD-fmk (20 μM) for 90 min. Immunoprecipitated endogenous FADD and total cell lysates were analysed by Western blotting. (**E**) Schematic representation of N4BP1 protein with indicated CASP8 cleavage sites. (**F**) The effect of N4BP1 and its cleavage fragments on cell viability. N4BP1^-/-^ MEFs reconstituted with empty vector, HA-N4BP1 (1-893), HA-N4BP1 (1-893, D311/484/488A), HA-N4BP1 (1-488) or HA-N4BP1 (489-893) were either exposed for 10 hours to TNFα (10 ng/ml) and CHX (0.5 μg/ml) or left untreated. Cell viability was determined by the CellTiter-Glo Luminescent assay. Three independent experimental replicates consisting of technical duplicates were performed. Results are shown as means and s.e.m. (n=3). n.s., no statistically significant difference, **** *P* < 0.0001, determined by two-way ANOVA, followed by *post hoc* Tukey’s multiple comparisons test. (**G**) Upon binding of TNFα, transmembrane TNFR1 undergoes trimerization, which enables formation of TNFR1-SC. Initially, TNFR1 trimer independently binds TRADD and RIP1. TRADD functions as a scaffold protein, which recruits Ub ligases cIAP1 and cIAP2 through TRAF2/5 to TNFR1-SC. cIAP1/2 attaches K63-linked Ub chains on RIP1 and automodifies itself, which is a prerequisite for LUBAC recruitment. Once recruited, LUBAC assembles M1 linkages on TNFR1-SC components, which are recognized by adapter subunit NEMO of IKK complex. Subsequently, LUBAC conjugates M1 linkages on NEMO, which is essential for the IKK activation. Next, IKK-mediated phosphorylation of NFκB inhibitor IºB leads to its K48 ubiquitination by SCF^β-TrCP^. Subsequent 26S proteasome-mediated removal of IºB enables nuclear translocation of NFºB/REL transcription factors and the induction of NFκB-dependent gene expression. We propose that N4BP1 regulates prosurvival TNFR1 signalling by recognizing linear Ub chains of nascent TNFR1-SC through its LUBIN. N4BP1 likely competes with linear Ub-specific DUBs (CYLD and OTULIN) for binding to M1 linkages, similarly to other linear Ub readers. During apoptosis, active CASP8 recognizes and cleaves N4BP1. N4BP1 (489-893) cleavage fragment, which contains LUBIN, undergoes proteasomal degradation. Consequently, the removal of C-terminal N4BP1 cleavage fragment promotes cell death.

Interestingly, we observed that prolonged TNFα stimulation results in the proteolytic cleavage of N4BP1 (**Extended Data Fig. 9A**). Important regulators of TNFR1 signalling, such as RIP1, CYLD and catalytic LUBAC subunit HOIP, are targets of CASP8-mediated cleavage^12,23,24^. Based on the computational analysis^25^, CASP8 was the most promising candidate to cleave N4BP1. Indeed, N4BP1 interacts and is processed by CASP8 both *in vivo* and *in vitro* (**Extended Data Fig. 9B–9E**), with cleavage pattern indicating multiple cleavage sites within N4BP1 (**Extended Data Fig. 9F–9G**). Mass spectrometry (MS)-based analysis of TNFα- and CHX-treated, immunoprecipitated N4BP1, identified two N4BP1 peptides ^477^QNSSCTVDLETD^488^ and ^297^QFSLENVPEGELLPD^311^ that are most likely a result of the CASP8 proteolytic activity (**Extended Data Fig. 10A–10B**). The mutational analysis confirmed N4BP1 residues D311 and D488 as major CASP8 recognition sites and showed that D484 residue is also cleaved, likely due to its close proximity to D488 (**Fig. 4E**, **Extended Data Fig. 10C**). Proteolytic processing of N4BP1 generates a N4BP1 (1-488) fragment, which preserves the ability to unspecifically bind Ub (**Extended Data Fig. 10D**), and the N4BP1 (489-893) fragment, which contains functional LUBIN (**Extended Data Fig. 10D**). N4BP1^-/-^ MEFs reconstituted with full-length HA-N4BP1 (1-893), uncleavable N4BP1 (1-893, D311/484/488A) or LUBIN-containing HA-N4BP1 (489-893) could suppress TNFα-mediated cell death, contrary to cells reconstituted with either empty vector or HA-N4BP1 (1-488) (**Fig. 4F**). The effect of LUBIN-containing N4BP1 fragment was striking, considering its very low expression level in N4BP1^-/-^ MEFs (**Extended Data Fig. 10E**). Noteworthy, we observed that LUBIN-containing N4BP1 fragment is unstable and undergoes proteasomal degradation **(Extended Data Fig. 10F)**. Hence, CASP8-mediated proteolytic processing of N4BP1 removes antiapoptotic LUBIN-containing fragment, facilitating apoptosis.

## Discussion

N4BP1 was initially described as a substrate of E3 ligase NEDD4^26^ and shown to inhibit ubiquitination and proteasomal degradation of E3 ligase ITCH substrates^27^. N4BP1 was also identified, but not further studied, as a Ub binder in a protein array and Ub-interactor affinity enrichment-MS (UbIA-MS) screens^18,28^. Nepravishta et al.,^15^ have recently described the structural model of CoCUN domain that encompasses the last 50 amino acids of human N4BP1 and binds to the I44 hydrophobic patch of monoUb, similar to our findings. Interestingly, related proteins MCPIP1-4 and KHNYN also contain divergent CUE domains (**Extended Fig. 1C**), but only KHNYN CUE binds Ub (data not shown and^29^).

MCPIP1 possesses ribonuclease activity within its PIN domain and plays a critical role in the inflammatory response by degrading mRNA of many cytokines^30^. MCPIP1 PIN domain undergoes head-to-tail intermolecular dimerization, enabling mRNA processing, and mutations preventing oligomerization abolish RNase activity^21^. N4BP1 was also recently identified as an interferon-inducible inhibitor of HIV-1 in primary T cells and macrophages, and shown to specifically degrade HIV-1 mRNA to control HIV-1 latency and reactivation^31^. Since RNase activity-deficient N4BP1 mutant failed to dimerize (data not shown), we were unable to determine the potential contribution of RNase activity to the effect of LUBIN in NFκB signalling. Future studies should systematically screen for specific N4BP1 RNase substrates and their potential role in NFκB signalling.

Two classes of structurally distinct LUBID interaction modes have been characterized so far^8^. The UBAN domains in NEMO, OPTN and ABIN form a parallel coiled-coil, which enables interactions with M1 linkages **(Extended Data Fig. 11A**). In contrast, HOIL-1L and A20 utilize structural elements that adopt a zinc finger fold to recognize and bind linear Ub chains with high affinity **(Extended Data Fig. 11B)**. Our findings indicate that N4BP1 utilizes a third and hitherto uncharacterized interaction mode, which depends on its RNase domain as a dimerization module **(Extended Data Fig. 11C)**. Homodimerization of N4BP1 elicits a specific orientation of the two CUE domains, which facilitates a formation of a highly stable complex with M1-linked diUb. Some CUE domains have been reported to self-associate and form functional dimers^32,33^. In fact, our analytical gel filtration experiments show that the elution volume of the CUE domain corresponds to a molecular weight of 8.4 kDa (**Extended Data Figs. 6C–6D**), which is larger than the calculated molecular weight (5.4 kDa) of a monomer, thus indicating a tendency for self-association that potentially contributes to linear chain specificity of the N4BP1 dimer. However, our ITC data of the isolated CUE domain do not show a marked preference to bind to linear Ub over monoUb (**Extended Data Fig. 2**). We therefore conclude that the potential of the CUE domain to dimerise is not sufficient to induce Ub chain binding specificity, which is only realised in the context of the N4BP1 constructs comprising the RNase domain. According to our knowledge, this is the first example of a UBD that does not display any Ub chain binding preference on its own, but can mediate a highly specific Ub chain interaction in combination with a dimerization module. Our results suggest a new combinatorial mechanism, which exploits the modular nature of domains with different functions to create linear Ub-binding specificity. Several LUBID-containing proteins also play an important role in TNFR1 signalling. UBAN-containing OPTN acts as a negative regulator of TNFR1 signalling upon TNFα stimulation, with linear Ub and CASP8 binding being critical for NFκB and apoptosis suppression respectively^34–36^. Abolished Ub binding by UBAN-containing protein ABIN-1 promotes NFκB and proinflammatory signalling, enhancing the production of proinflammatory mediators^37–39^. Furthermore, ABIN-1 prevents cell death by inhibiting CASP8 recruitment to FADD in UBAN-dependent manner^40^, presumably by binding linear Ub-modified FADD that prevents it from forming complex II with CASP8^12^. Ub-editing enzyme A20 is recruited to TNFR1-SC through its M1-(ZnF7) and K63-specific (ZnF4) UBDs^41,42^. By protecting linear Ub chains from DUB cleavage, A20 prevents formation of the complex II^42^. These data are in agreement with our results, showing how distinct linear Ub readers within TNFR1-SC regulate TNFR1 signalling, as well as with data showing how deficiency of LUBAC components promotes complex II assembly and induces aberrant TNF-mediated endothelial cell death^43^.

Gitlin et al.^1^ identified N4BP1 as a suppressor of cytokine production that is inactivated by CASP8, similar to our findings. Mice lacking N4BP1 show increased production of a subset of cytokines, which is in agreement with our data. We here provide a detailed mechanism on how N4BP1 exerts its activity in TNFR1 signalling through the newly identified LUBIN, and explain how CASP8 cleavage leads to removal of functional LUBIN that is no longer able to stabilize linear Ub linkages and the integrity of TNFR1-SC. Similarly, HOIP is also cleaved upon the induction of apoptosis and subjected to proteasomal degradation. Whereas the C-terminal fragment of HOIP retains NFκB activity, linear ubiquitination of NEMO and FADD are decreased^12^, facilitating the assembly of complex II and apoptosis. It was also recently shown that CASP8 inhibitor cFLIP is modified by LUBAC to prevent its proteasomal degradation. Inactivation or depletion of HOIP leads to the removal of cFLIP, releasing CASP8 inhibition^44^. This implies that multiple proapoptotic regulators are kept in check by linear Ub chains generated by LUBAC and that the removal of these linkages unleashes apoptosis.

In summary, we propose that N4BP1 is recruited to the nascent TNFR1-SC upon TNFα stimulation, where it binds linear Ub linkages via LUBIN to regulate the integrity of TNFR1-SC and duration of proinflammatory signalling (**Fig. 4G**). Prolonged TNFα stimulation leads to the activation of CASP8 that cleaves N4BP1. After the C-terminal part of N4BP1 is removed by 26S proteasome, complex II formation and apoptosis are accelerated. Thus, N4BP1 utilizes a complex mechanism to regulate the progression of the TNFα-mediated NFκB pathway and cell death.

## Acknowledgments

We thank Michael Kuehn for N4BP1^-/-^ MEFs. We are grateful to John Blenis, Irmela Jeremias and Clarissa von Haefen for providing CASP8-deficient Jurkat cells. We thank Tanya Sultana Masood for helping with protein purifications. We thank Jean Berthelet, Krishnaraj Rajalingam and Simin Rahighi for initial help with the project. We thank Ulrich Maurer and Lina Katharina Schlicher for helpful advice with the TNFα IP experiment. We thank Jaime Lopez-Mosqueda, Gergely Imre, Huib Ovaa and Andrea Gubas for essential discussions, comments and critical reading of manuscript. We also thank Ivan Dikic, Errol Friedberg, Caixia Guo, Michael Kuehn, Rodolfo Murillas and Adrian Ting for providing reagents. This work was supported by the Francis Crick Institute through provision of access to the MRC Biomedical NMR Centre. The Francis Crick Institute receives its core funding from Cancer Research UK (FC001029), the UK Medical Research Council (FC001029), and the Wellcome Trust (FC001029). KK was supported by the UPStream grant (EU, FP7, ITN project 290257).

## Author contributions

KoH, KK and BS conceived the study and wrote the manuscript with contributions from the remaining authors. KoH, KK, WS and BS designed, performed and analysed experiments. SS, SK, IP, DK, JG, AF and AP performed and analysed experiments.

## Data and material availability

All the plasmids generated in this study will be available upon request. All data is available in the main text or the supplementary materials.

## Supplementary Materials

Materials and Methods

Extended Discussion

Supplementary tables 1-4

Supplementary table S5 (separate Excel sheet)

Extended Data Figures 1 – 11

Extended Data Figure 1 – 11 Legends

References (1-23)

---

**Extended Data Figure 1.**
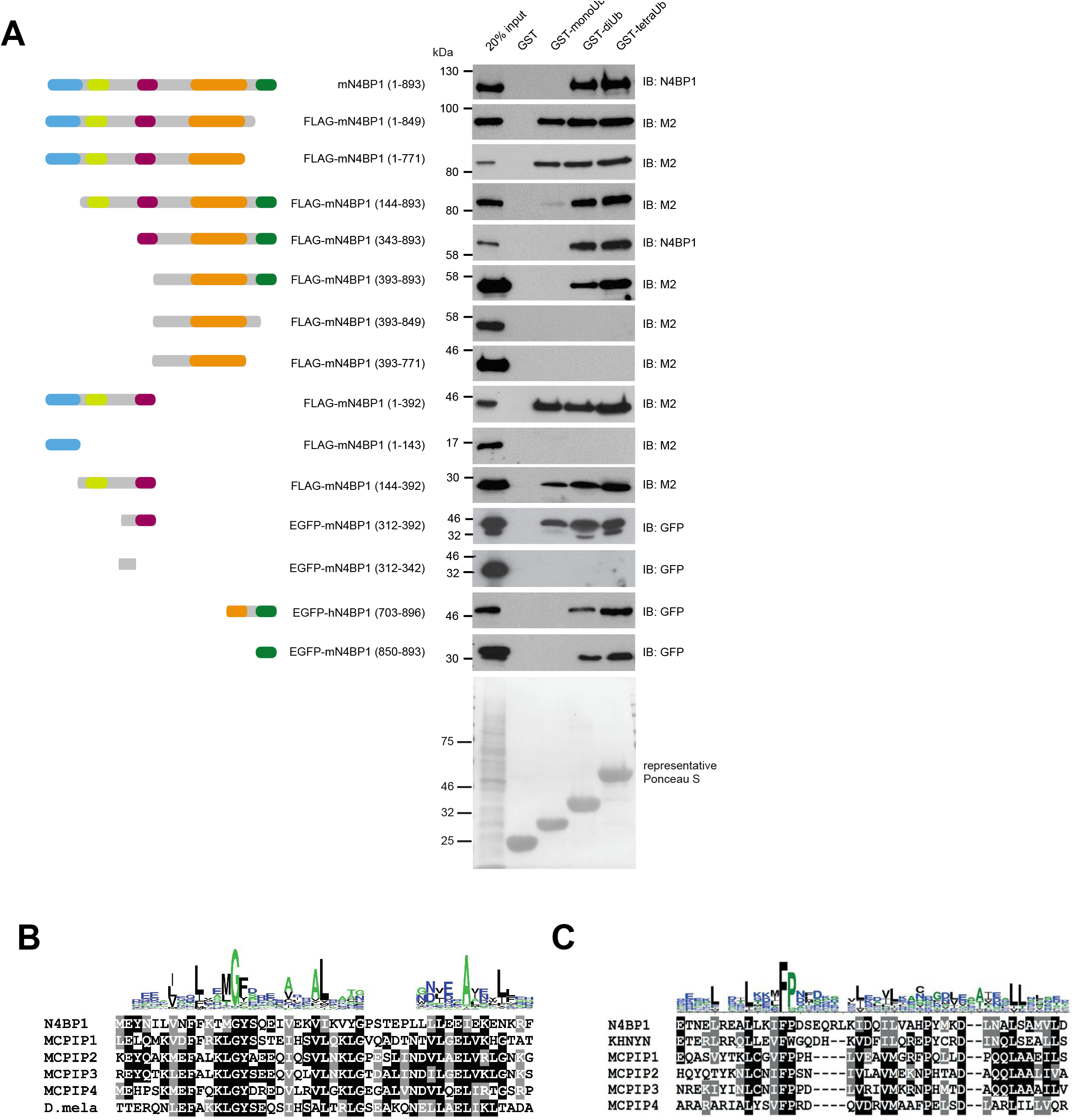

**Extended Data Figure 2.**
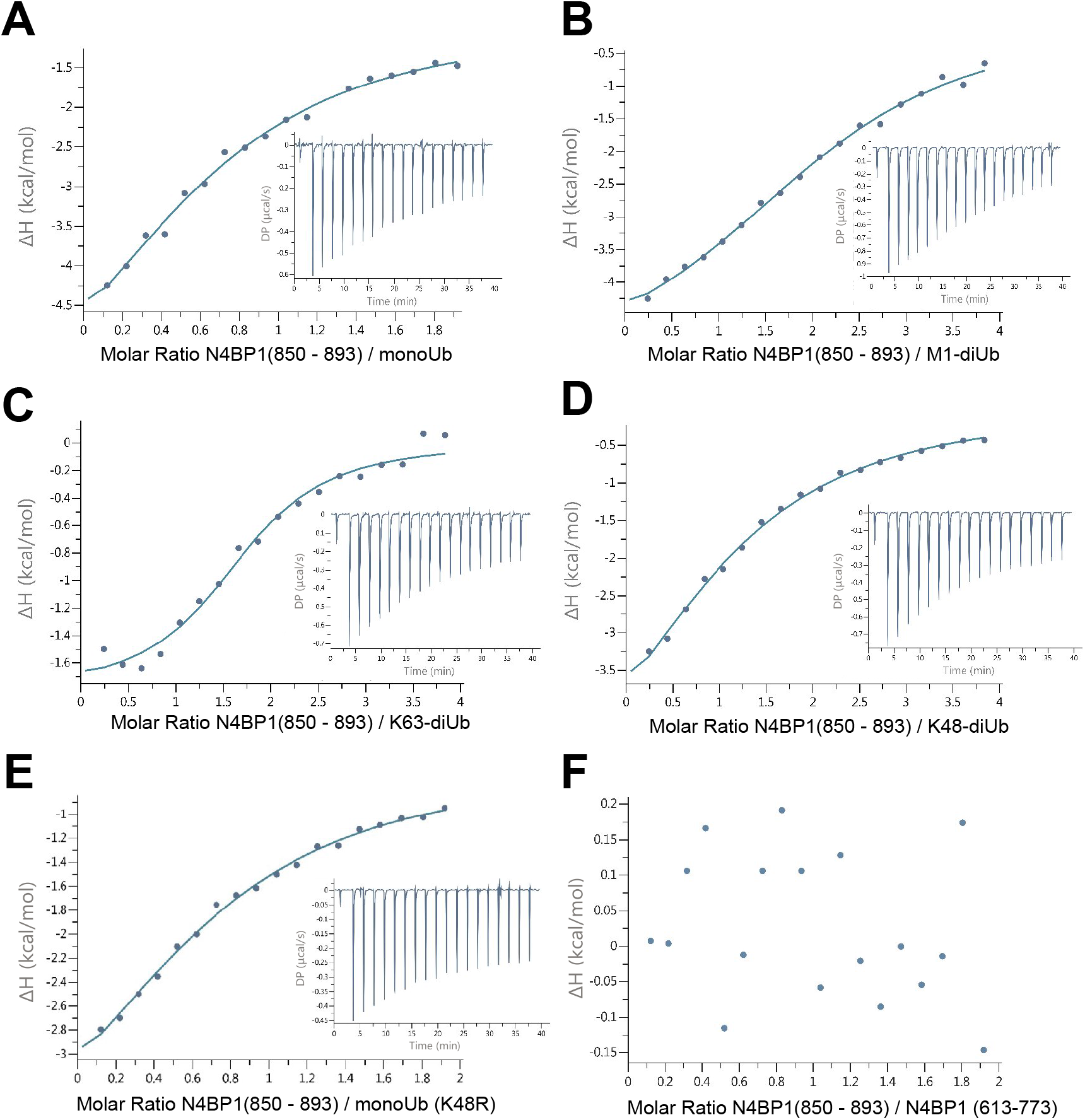

**Extended Data Figure 3.**
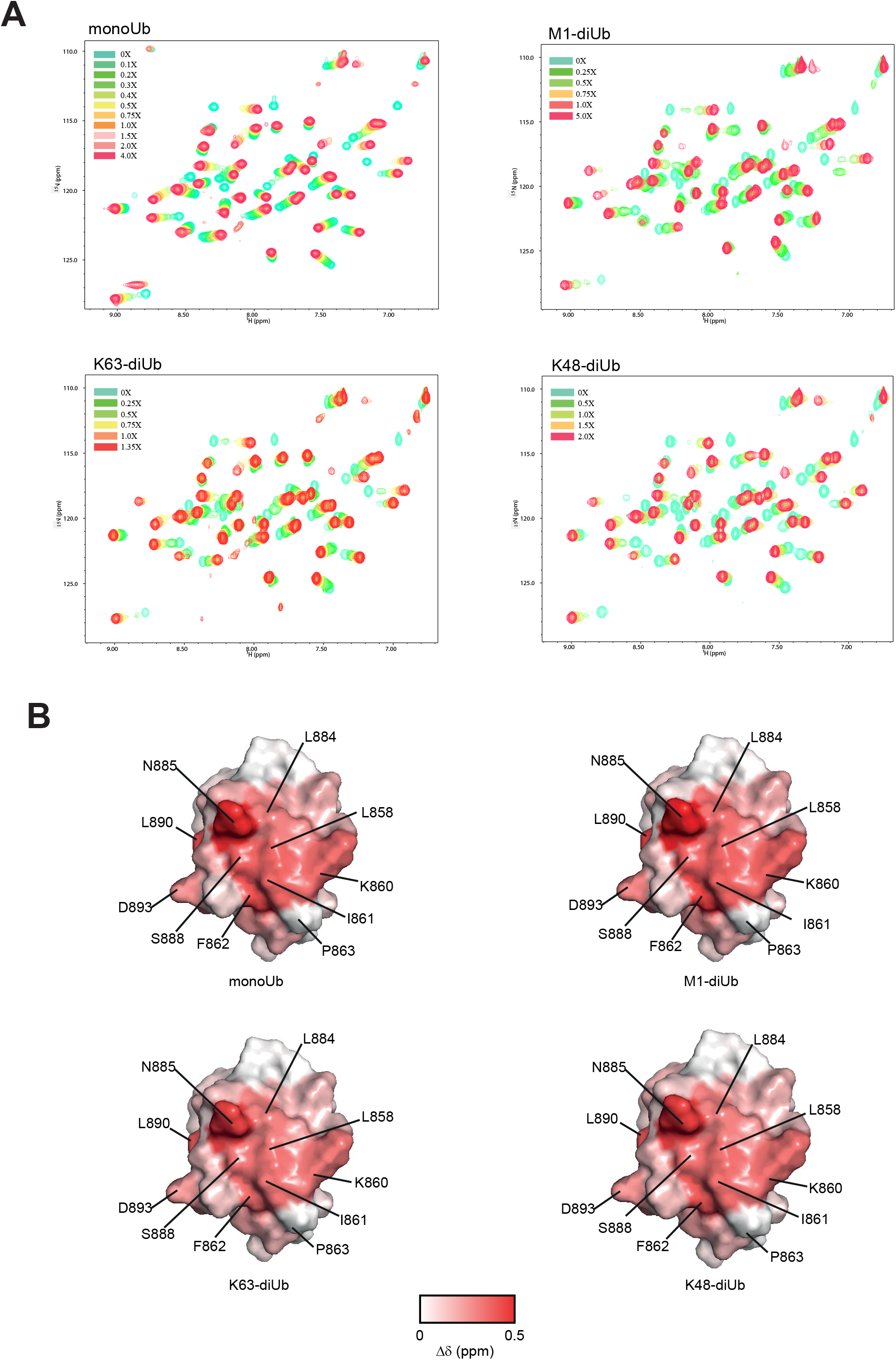

**Extended Data Figure 4.**
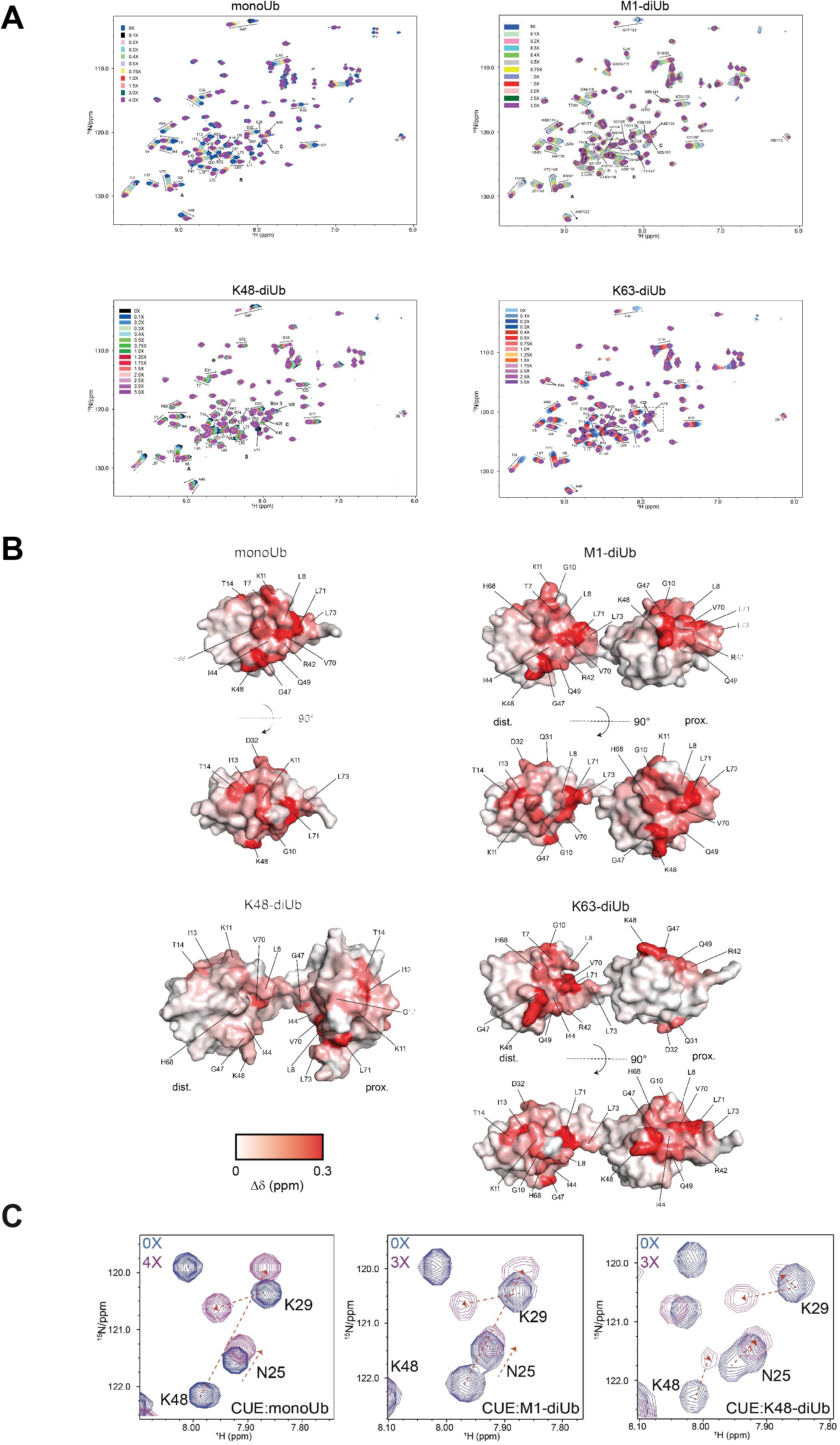

**Extended Data Figure 5.**
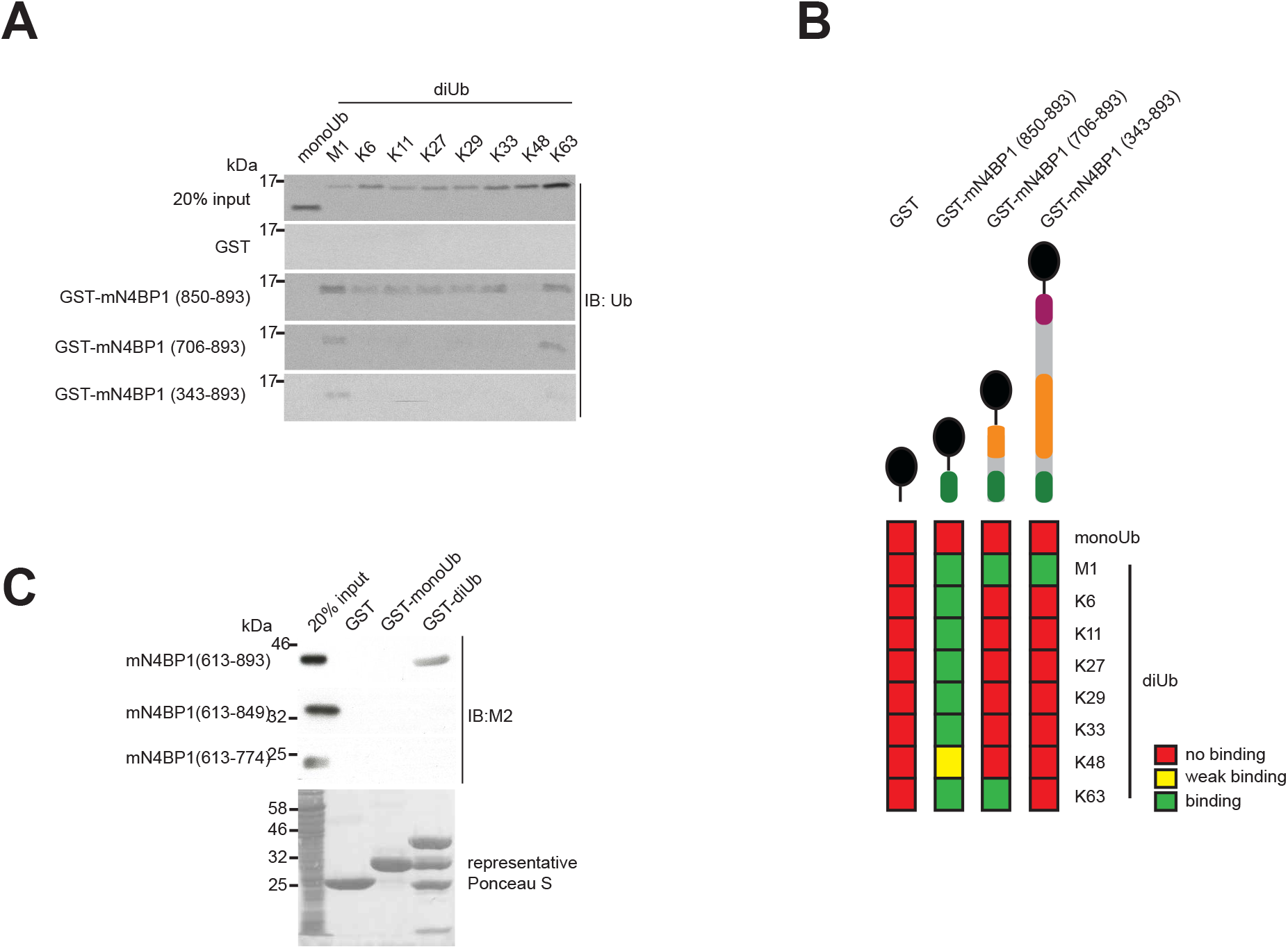

**Extended Data Figure 6.**
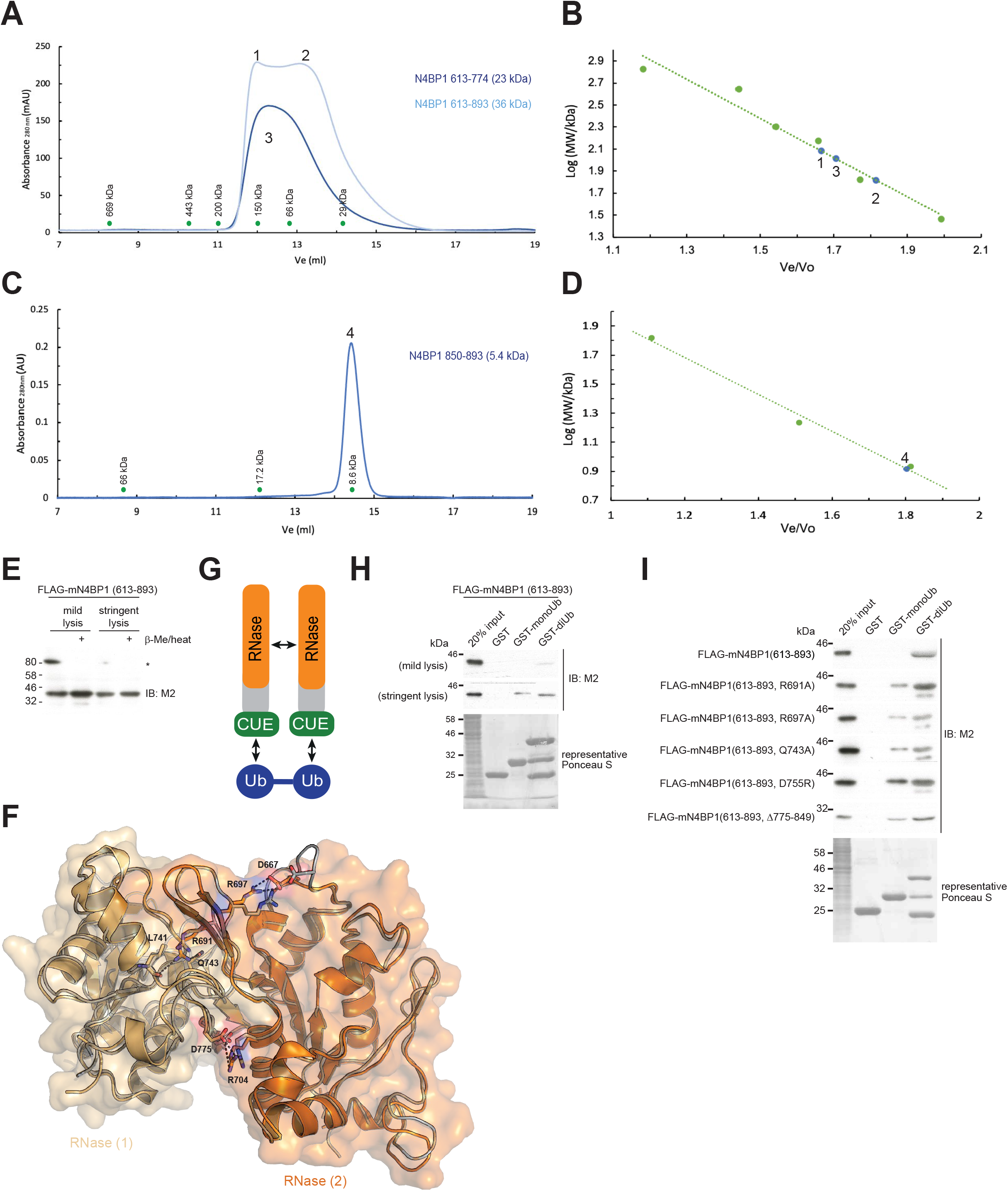

**Extended Data Figure 7.**
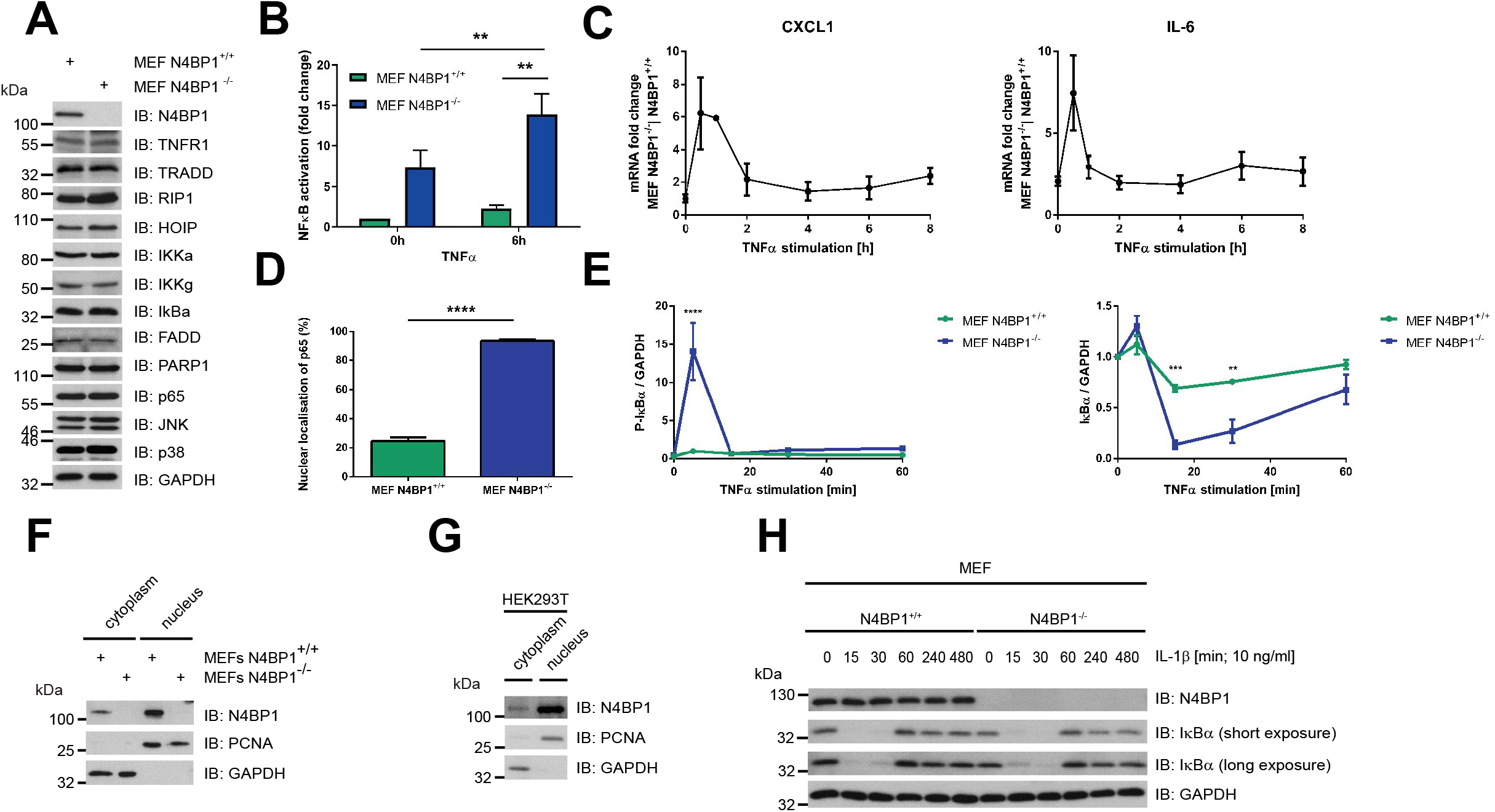

**Extended Data Figure 8.**
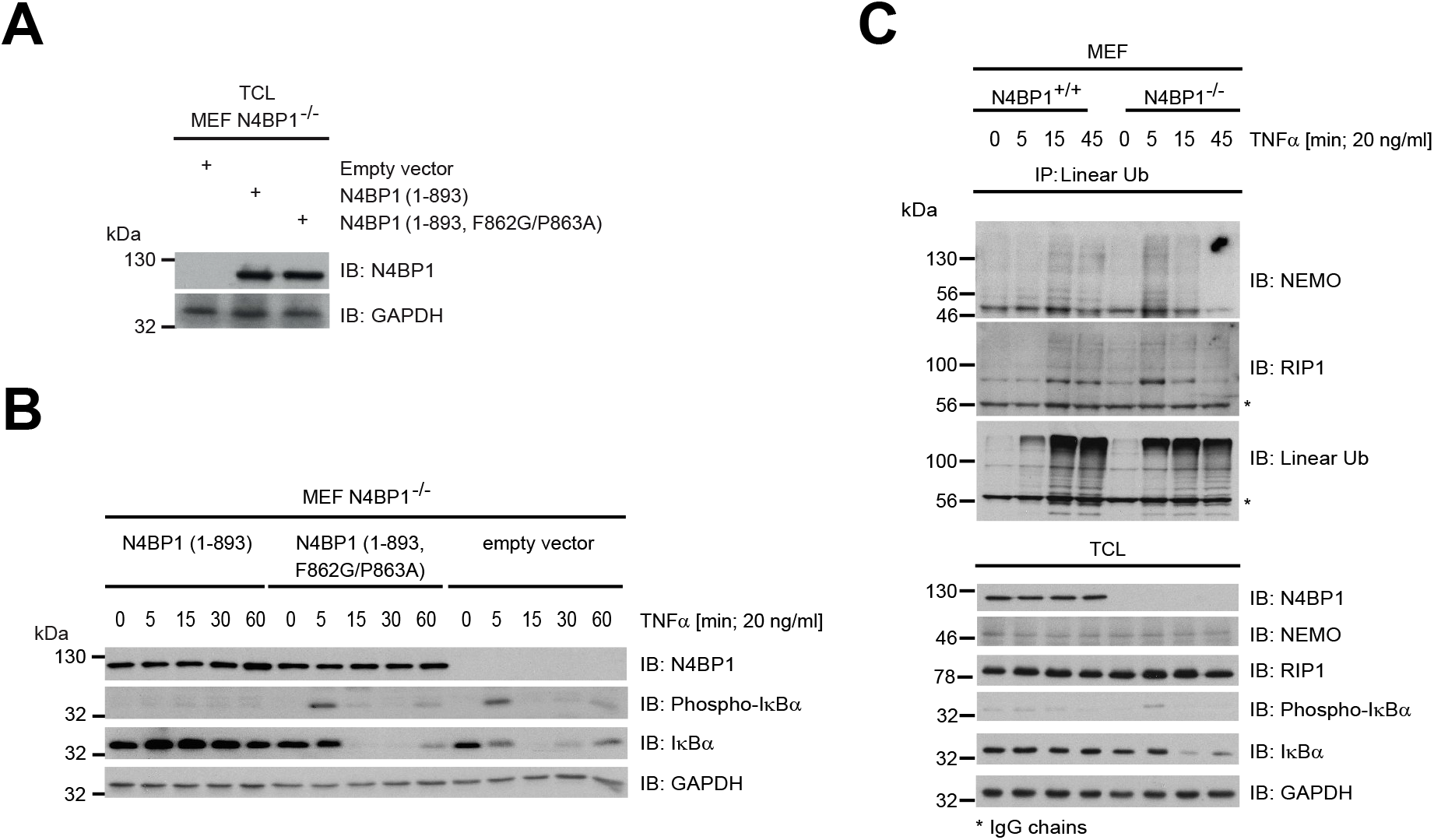

**Extended Data Figure 9.**
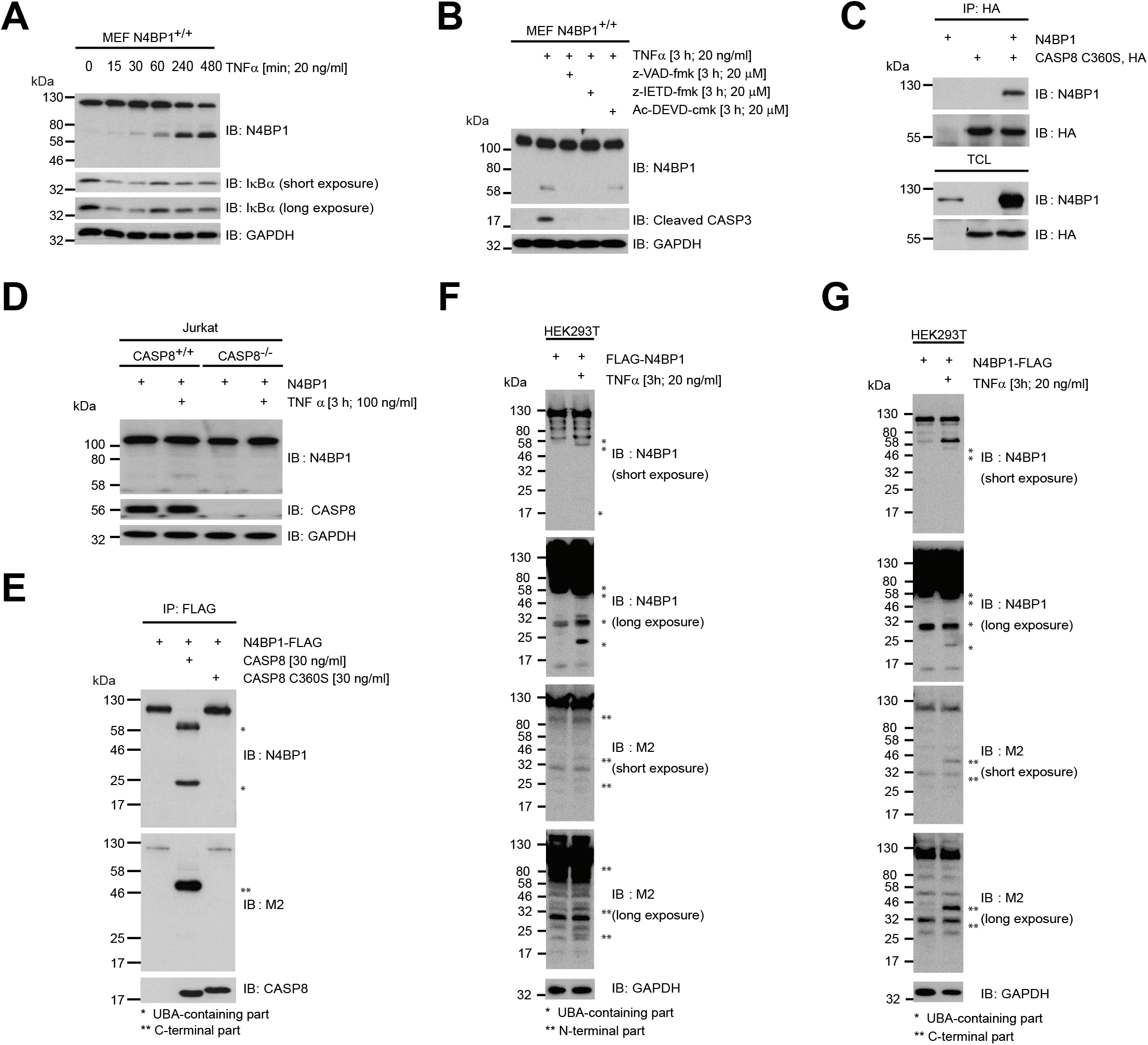

**Extended Data Figure 10.**
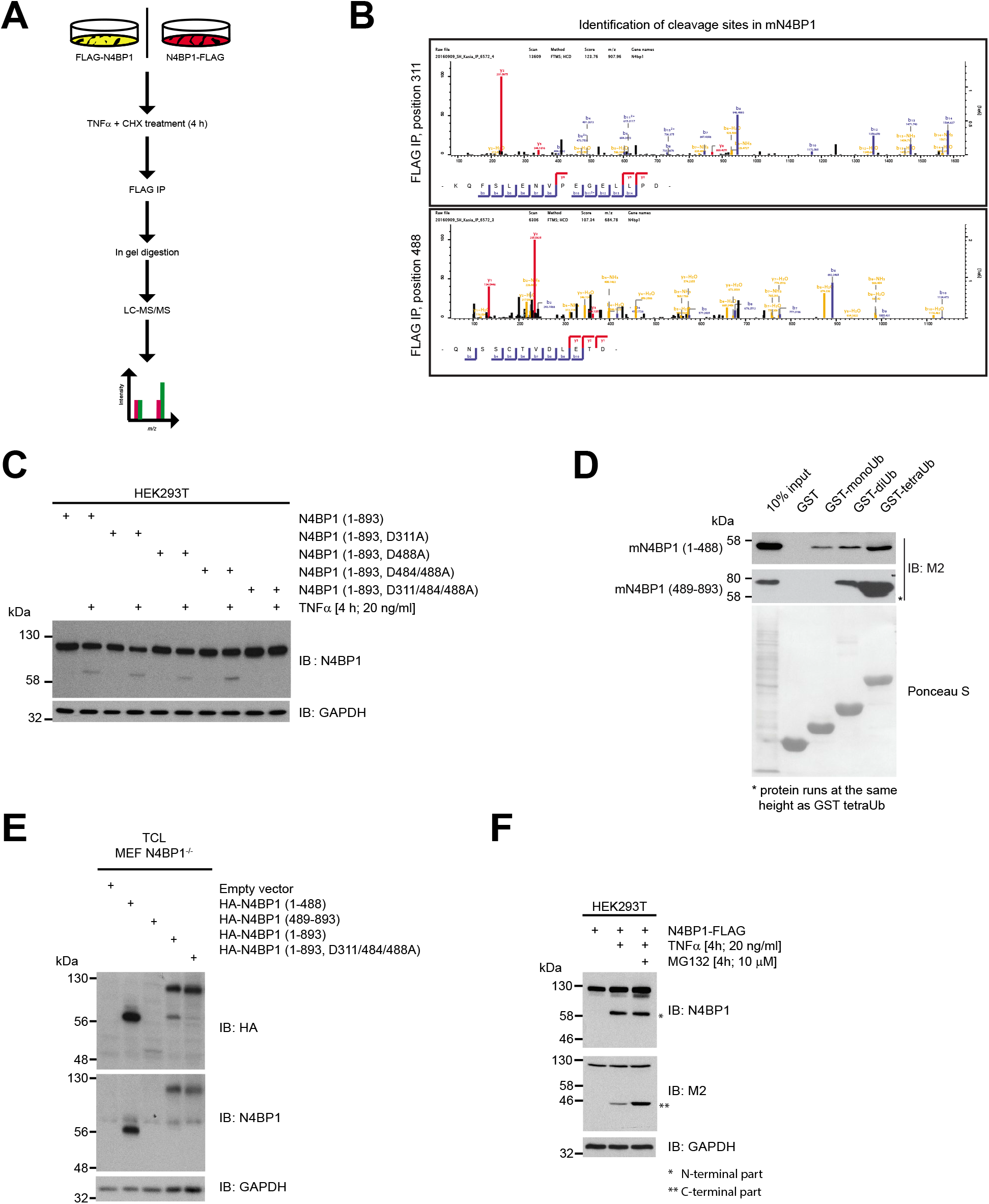

**Extended Data Figure 11.**
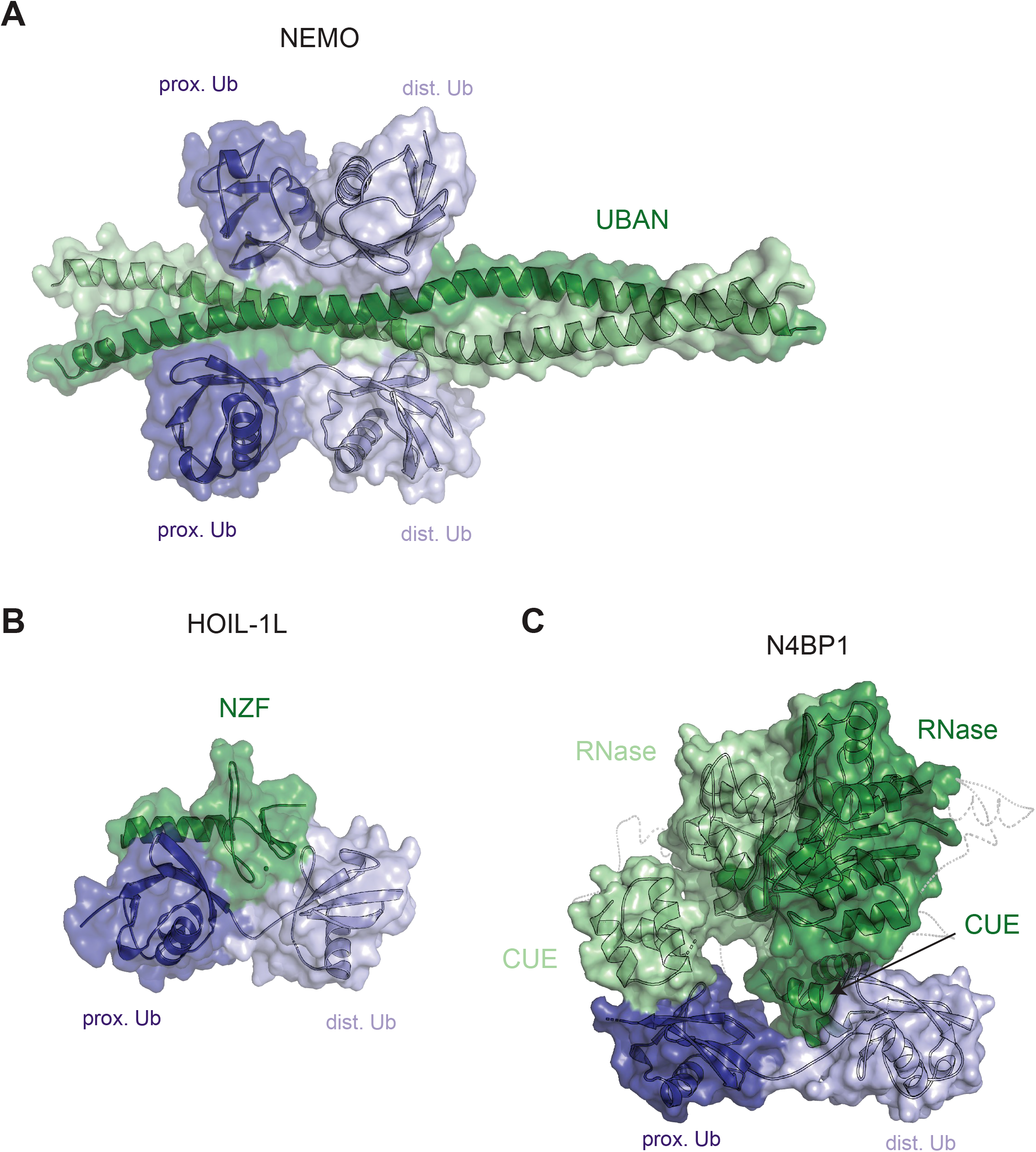

## References

1. Gitlin, A. D. et al. Integration of innate immune signalling by caspase-8 cleavage of N4BP1. Nature 587, 275–280 (2020).

2. Walczak, H. TNF and ubiquitin at the crossroads of gene activation, cell death, inflammation, and cancer. Immunological Reviews vol. 244 9–28 (2011).

3. Haas, T. L. et al. Recruitment of the Linear Ubiquitin Chain Assembly Complex Stabilizes the TNF-R1 Signaling Complex and Is Required for TNF-Mediated Gene Induction. Mol. Cell 36, 831–844 (2009).

4. Wertz, I. E. et al. Phosphorylation and linear ubiquitin direct A20 inhibition of inflammation. Nature 528, 370–375 (2015).

5. Kirisako, T. et al. A ubiquitin ligase complex assembles linear polyubiquitin chains. EMBO J. 25, 4877–4887 (2006).

6. Emmerich, C. H. et al. Lys63/Met1-hybrid ubiquitin chains are commonly formed during the activation of innate immune signalling. Biochem. Biophys. Res. Commun. 474, 452–461 (2016).

7. Gerlach, B. et al. Linear ubiquitination prevents inflammation and regulates immune signalling. Nature 471, 591–596 (2011).

8. Fennell, L. M., Rahighi, S. & Ikeda, F. Linear ubiquitin chain-binding domains. FEBS Journal vol. 285 2746–2761 (2018).

9. Dynek, J. N. et al. C-IAP1 and UbcH5 promote K11-linked polyubiquitination of RIP1 in TNF signalling. EMBO J. 29, 4198–4209 (2010).

10. Rahighi, S. et al. Specific Recognition of Linear Ubiquitin Chains by NEMO Is Important for NF-κB Activation. Cell 136, 1098–1109 (2009).

11. Webster, J. D. & Vucic, D. The Balance of TNF Mediated Pathways Regulates Inflammatory Cell Death Signaling in Healthy and Diseased Tissues. Frontiers in Cell and Developmental Biology vol. 8 (2020).

12. Goto, E. & Tokunaga, F. Decreased linear ubiquitination of NEMO and FADD on apoptosis with caspase-mediated cleavage of HOIP. Biochem. Biophys. Res. Commun. 485, 152–159 (2017).

13. Peltzer, N. et al. LUBAC is essential for embryogenesis by preventing cell death and enabling haematopoiesis. Nature 557, 112–117 (2018).

14. Kang, R. S. et al. Solution structure of a CUE-ubiquitin complex reveals a conserved mode of ubiquitin binding. Cell 113, 621–630 (2003).

15. Nepravishta, R. et al. CoCUN, a novel ubiquitin binding domain identified in N4BP1. Biomolecules 9, (2019).

16. Shih, S. C. et al. A ubiquitin-binding motif required for intramolecular monoubiquitylation, the CUE domain. EMBO J. 22, 1273–1281 (2003).

17. Hurley, J. H., Lee, S. & Prag, G. Ubiquitin-binding domains. Biochemical Journal vol. 399 361–372 (2006).

18. Zhang, X. et al. An Interaction Landscape of Ubiquitin Signaling. Mol. Cell 65, 941–955.e8 (2017).

19. Faesen, A. C. et al. The differential modulation of USP activity by internal regulatory domains, interactors and eight ubiquitin chain types. Chem. Biol. 18, 1550–1561 (2011).

20. Licchesi, J. D. F. et al. An ankyrin-repeat ubiquitin-binding domain determines TRABID’s specificity for atypical ubiquitin chains. Nat. Struct. Mol. Biol. 19, 62–72 (2012).

21. Yokogawa, M. et al. Structural basis for the regulation of enzymatic activity of Regnase-1 by domain-domain interactions. Sci. Rep. (2016) doi:10.1038/srep22324.

22. Itzhak, D. N., Tyanova, S., Cox, J. & Borner, G. H. H. Global, quantitative and dynamic mapping of protein subcellular localization. Elife 5, (2016).

23. O’Donnell, M. A. et al. Caspase 8 inhibits programmed necrosis by processing CYLD. Nat. Cell Biol. 13, 1437–1442 (2011).

24. Zhang, L. et al. RIP1 Cleavage in the Kinase Domain Regulates TRAIL-Induced NF-κB Activation and Lymphoma Survival. Mol. Cell. Biol. (2015) doi:10.1128/mcb.00692-15.

25. Gasteiger, E. et al. Protein Identification and Analysis Tools on the ExPASy Server. in The Proteomics Protocols Handbook 571–607 (2005). doi:10.1385/1-59259-890-0:571.

26. Murillas, R., Simms, K. S., Hatakeyama, S., Weissman, A. M. & Kuehn, M. R. Identification of developmentally expressed proteins that functionally interact with Nedd4 ubiquitin ligase. J. Biol. Chem. 277, 2897–2907 (2002).

27. Oberst, A. et al. The Nedd4-binding partner 1 (N4BP1) protein is an inhibitor of the E3 ligase Itch. Proc. Natl. Acad. Sci. U. S. A. 104, 11280–11285 (2007).

28. Fenner, B. J., Scannell, M. & Prehn, J. H. M. Identification of polyubiquitin binding proteins involved in NT-i<B signaling using protein arrays. Biochim. Biophys. Acta – Proteins Proteomics 1794, 1010–1016 (2009).

29. Castagnoli, L. et al. Selectivity of the CUBAN domain in the recognition of ubiquitin and NEDD8. FEBS J. 286, 653–677 (2019).

30. Musson, R., Szukała, W. & Jura, J. Mcpip1 rnase and its multifaceted role. International Journal of Molecular Sciences vol. 21 1–14 (2020).

31. Yamasoba, D. et al. N4BP1 restricts HIV-1 and its inactivation by MALT1 promotes viral reactivation. Nat. Microbiol. 4, 1532–1544 (2019).

32. Mitra, S., Traughber, C. A., Brannon, M. K., Gomez, S. & Capelluto, D. G. S. Ubiquitin interacts with the tollip C2 and CUE domains and inhibits binding of tollip to phosphoinositides. J. Biol. Chem. 288, 25780–25791 (2013).

33. G, P. et al. Mechanism of ubiquitin recognition by the CUE domain of Vps9p. Cell 113, 609–620 (2003).

34. Zhu, G., Wu, C. J., Zhao, Y. & Ashwell, J. D. Optineurin Negatively Regulates TNFα-Induced NF-κB Activation by Competing with NEMO for Ubiquitinated RIP. Curr. Biol. (2007) doi:10.1016/j.cub.2007.07.041.

35. Sudhakar, C., Nagabhushana, A., Jain, N. & Swarup, G. NF-κB mediates tumor necrosis factor α-induced expression of optineurin, a negative regulator of NF-κB. PLoS One 4, (2009).

36. Nakazawa, S. et al. Linear ubiquitination is involved in the pathogenesis of optineurin-associated amyotrophic lateral sclerosis. Nat. Commun. 7, (2016).

37. Nanda, S. K. et al. Polyubiquitin binding to ABIN1 is required to prevent autoimmunity. J. Exp. Med. 208, 1215–1228 (2011).

38. Heyninck, K., Kreike, M. M. & Beyaert, R. Structure-function analysis of the A20-binding inhibitor of NF-κB activation, ABIN-1. FEBS Lett. 536, 135–140 (2003).

39. Wagner, S. et al. Ubiquitin binding mediates the NF-κB inhibitory potential of ABIN proteins. Oncogene 27, 3739–3745 (2008).

40. Oshima, S. et al. ABIN-1 is a ubiquitin sensor that restricts cell death and sustains embryonic development. Nature 457, 906–909 (2009).

41. Razani, B. et al. Non-catalytic ubiquitin binding by A20 prevents psoriatic arthritis-like disease and inflammation. Nat. Immunol. 21, 422–433 (2020).

42. Martens, A. et al. Two distinct ubiquitin-binding motifs in A20 mediate its anti-inflammatory and cell-protective activities. Nat. Immunol. 21, 381–387 (2020).

43. Peltzer, N. et al. HOIP deficiency causes embryonic lethality by aberrant TNFR1-mediated endothelial cell death. Cell Rep. 9, 153–165 (2014).

44. Tang, Y. et al. Linear ubiquitination of cFLIP induced by LUBAC contributes to TNF-induced apoptosis. J. Biol. Chem. 293, 20062–20072 (2018).

