## Supplementary text for "N4BP1 is dimerization-dependent linear ubiquitin reader regulating TNFR1 signalling through linear ubiquitin binding and Caspase-8-mediated processing"

^1^Institute of Biochemistry II, Goethe University School of Medicine, Frankfurt (Main), Germany. ^2^School of Biological and Behavioural Sciences, Queen Mary University of London, London, United Kingdom. ^3^Structural Biology Science Technology Platform, Francis Crick Institute, London, United Kingdom. ^4^School of Physical and Chemical Sciences, Queen Mary University of London, London, United Kingdom. ^5^Department of Computer Science, Brunel University London, Uxbridge, United Kingdom. ^6^Institute for Genetics, University of Cologne, Cologne, Germany. ^7^Centre for Host-Microbiome Interactions, Dental Institute, King’s College London, London, United Kingdom. ^8^Current address: Department of Molecular Biology, Radboud Institute for Molecular Life Sciences, Radboud University, Nijmegen, The Netherlands. ^9^Current address: Department of Oncology, University of Oxford, Oxford, United Kingdom. ^10^Current address: Department of Biology, University of York, Wentworth Way, York, United Kingdom.^11^These authors contributed equally. ^12^Correspondence: [](about:blank), [](about:blank), [](about:blank).

**This PDF file includes:**

Materials and Methods

Extended discussion

Extended Data Figure 1-11 Legends

Supplementary tables 1-4

References (1-23)

**Other Supplementary Materials for this manuscript include the following:**

Supplementary table S5 (related to **Extended Figure 10B**) with mass spectrometry data supporting the identification of N4BP1 cleavage sites.

**EXPERIMENTAL MODEL AND SUBJECT DETAILS**

**Cell lines**

Cell line HEK293T was purchased from ATCC. Wild-type and CASP8-deficient Jurkat cells were generated by John Blenis (USA) and Clarissa von Haefen (Germany). Primary N4BP1^+/+^ and N4BP1^-/-^ MEFs, kindly provided by Michael Kuehn (USA), were immortalized by transfection with plasmid containing SV40 T-antigen. All cells were regularly checked for *Mycoplasma* infection using VenorGeM Classic from *Minerva Biolabs GmbH* (Ltd).

**Reagents**

Puromycin and zeocin were from *Invivogen*. Ac-DEVD-cmk, z-IETD-fmk and z-VAD-fmk were from *Santa Cruz Biotechnology*. TNFα and IL-1β were from *PeproTech*. Polybrene and benzonase endonuclease were from *Merck*. Phosphatase and Protease Inhibitor Cocktails were from *Roche Applied Science*. Crystal violet was from *Carl Roth GmbH* and cycloheximide (CHX) from *Enzo* *Life Sciences*.

**Plasmids and antibodies**

The lists of used plasmids and antibodies are available in **Supplementary tables 1** and **2**, respectively. In hexaUb YTH9 plasmid, five tandem Ub molecules had a terminal GG motif mutated to GV (to prevent cleavage by cellular DUBs during the screen), whereas the proximal Ub lacked GG motif (to prevent its potential conjugation to yeast proteins). For SPR experiments, the N4BP1 (613-774) and N4BP1 (613-893) fragments were cloned into pCold-TF (*Takara*) vector. For analytical gel filtration, the N4BP1 (613-774) and N4BP1 (613-893) fragments were cloned into pCold-I (*Takara*) vector, which include an N-terminal 20 amino acid solubility tag (GGGTPKAPNLEPPLPEEEKEG).

**METHOD DETAILS**

**Yeast two-hybrid**

Gal4BD-fused hexaUb plasmid was transformed into Y2HGold *Saccharomyces cerevisiae* strain (*Clontech*) and used as bait in Y2H screen, where it was mated with the human normalized cDNA library (*Clontech*) transformed into Y187 yeast strain (prey). Four independent reporter genes (*AUR1-C*, *ADE2*, *HIS3*, and *MEL1*) were used for selection according to the manufacturer's instructions. Clone identities were determined by Sanger sequencing (*Microsynth Seqlab*).

**Bioinformatics analysis**

Multiple alignments were calculated using the L-Ins-I algorithm of the MAFFT package^1^. Sequence database searches were performed by the generalized profile method^2^, using the pftools package. Inter-family similarities were established by Hidden Markov Model (HMM) to HMM comparisons, using the HHSEARCH program package^3^.

**Transfection of mammalian cells**

HEK293T cells were transiently transfected with either polyethylenimine (PEI, *Polysciences*) or GeneJuice transfection reagent (*Merck Millipore*). Non-adherent Jurkat cells were electroporated using the Neon Transfection System (*Invitrogen*). MEFs were transiently transfected using GenJet *In Vitro* DNA Transfection Reagent (*SignaGen Laboratories*). All the transfections were performed according to the manufacturer's instructions. If the experiment required transfection of several plasmids simultaneously, appropriate amounts of empty vectors were used to equalize the total plasmid amount across the samples.

**Retroviral production**

For reconstitution of N4BP1^-/-^ MEFs, various pBabe plasmids (carrying resistance to either puromycin or zeocin) were used. Twenty-four hours after seeding, HEK293T cells were transfected with appropriate plasmid DNA and helper plasmid phai by using GeneJuice. Thirty six hours post-transfection, DMEM containing retroviruses was filtered, mixed with polybrene (final concentration 4 μg/ml) and transferred to target cells. Forty-eight hours post-infection, selection with either 300 μg/ml of zeocin or 4 μg/ml of puromycin was started. Cells were cultured in the presence of appropriate antibiotics throughout use.

**Various treatments of mammalian cells**

Where indicated, starved cells were treated with either recombinant human IL-1β, mouse TNFα or human HS-TNFα at concentrations 10 ng/ml, 20 ng/ml and 1 μg/ml, respectively. To identify N4BP1 cleavage sites, serum-starved HEK293T cells were treated with recombinant mouse TNFα (20 ng/ml) and additionally, with proteasome inhibitor MG132 (*Tocris Bioscience*) at 10 μM final concentration. For identification of the specific CASP responsible for N4BP1 proteolysis, MEF N4BP1^+/+^ and HEK293T cells were treated with recombinant mouse TNFα (20 ng/ml) and 20 μM of either CASP inhibitors z-VAD-fmk, z-IETD-fmk or Ac-DEVD-cmk. For testing N4BP1 proteolysis in Jurkat cells, TNFα was used at concentration 100 ng/ml. For cell death induction, recombinant mouse TNFα and cycloheximide (CHX) were used at concentration 10 ng/ml and 0.5-1 μg/ml, respectively. For inhibition of apoptosis, cells were treated with mouse TNFα (10 ng/ml), CHX (1 μg/ml) and z-VAD-fmk (20 μM). The duration of treatments is indicated in the figures.

**Expression and purification of recombinant proteins**

Various GST fusions were purified as shown previously^4^. HS-TNFα was expressed in *E. coli* with an N-terminal STREP II tag and purified on Strep-tactin Sepharose according to manufacturer’s instructions (*GE Healthcare*), followed by gel filtration using Superdex S75 (*GE Healthcare*) and subsequent dialysis against PBS. Recombinant active and inactive (C360S) CASP8 were purified from BL21 *E. coli* by lysing bacterial pellet in 50 mM Na_2_HPO_4_/NaH_2_PO_4_ (pH 7.4), 300 mM NaCl, 20 mM imidazole, 30% glycerol and 20 mM β-mercaptoethanol (β-Me). Precleared lysates were bound to nickel resins (*Qiagen*) and washed several times with lysis buffer, after which proteins were eluted with 50 mM Na_2_HPO_4_/NaH_2_PO_4_ (pH 8.0), 300 mM NaCl, 250 mM imidazole, 30% glycerol and 20 mM β-Me. N4BP1 proteins for SPR and ITC measurements were expressed and purified from BL21 *E. coli* using GST or immobilized cobalt affinity chromatography followed by size exclusion chromatography and anion exchange chromatography, if required. Size exclusion chromatography was performed in 50 mM HEPES (pH 7.4), 150 mM NaCl and 1 mM DTT, using Superdex S200 or Superdex S75 chromatography columns *(GE Healthcare)*. For anion exchange chromatography, protein samples were separated by varied gradient elution from MonoQ columns *(GE Healthcare)* using 20 mM HEPES (pH 8.5), 1 mM DTT, 0.5% Triton X-100 and 0.1-1.0 M NaCl. For NMR spectroscopy ^15^N/^13^C-labelled N4BP1 (850-893) was expressed in M9 media (6 g/L Na_2_HPO_4_, 3 g/L KH_2_PO_4_, 0.5 g/L NaCl, 0.7 g/L ^15^NH_4_Cl, 2 g/L ^13^C-D-glucose, 10 mL/L 100X MEM vitamin solution *(Gibco)*, 10 μM FeSO_4_, 10 μM CaCl_2_, 2 mM MgSO_4_, pH 7.4). For ITC and NMR experiments, the His-tag was removed from isolated N4BP1 CUE domain by incubation with 3C protease at 4 °C, overnight. For analytical gel filtration, the His-tag of purified N4BP1 fragments was not removed. Enzymatic synthesis and purification of K48- and K63-linked Ub was essentially carried out as described^5^.

**Analytical size exclusion chromatography**

Analytical size exclusion chromatography was performed using a Superose^TM^ 12 10/300L column (*GE Healthcare*) calibrated with the 29,000-700,000 Da GF Marker Kit (*Sigma-Aldrich*). Protein samples were eluted in 50 mM HEPES, pH 8.5, 150 mM NaCl, 1 mM DTT, 0.5% Triton X-100.

**Isothermal titration calorimetry (ITC)**

ITC experiments were performed at 293 K using a Microcal PEAQ-ITC calorimeter (*Malvern*). The protein solutions were prepared in a buffer containing 50 mM HEPES (pH 7.4), 50 mM NaCl and 0.5 mM TCEP. Experiments were performed at cell at concentrations 50-100 μM. The injectant concentration in the syringe was usually 10 fold to the titrant. For each titration 20 injections of 2 μl were performed. Integrated data, corrected for heats of dilution, were fitted using a nonlinear least-squares algorithm to obtain a binding curve, using the MicroCal Origin 7.0 software package. Each experiment was repeated at least twice, and average values are reported in **Table 1**.

***In vitro* CASP8 cleavage assay**

C-terminally FLAG-tagged N4BP1 was transiently transfected into HEK293T cells, which were washed in 1xPBS 24 hours after transfection. Cells were lysed in 1% Triton X-100 buffer (50 mM Tris-HCl, pH 7.5; 40 mM NaCl; 5 mM EDTA; 1% Triton X-100; 1x Protease inhibitor cocktail) and incubated on ice for 20 min. Precleared lysates were incubated with M2 agarose (*Sigma-Aldrich*) at 4°C for 2 h. After 5 washing steps with modified 1% Triton X-100 buffer (50 mM Tris-HCl, pH 7.5; 500 mM NaCl; 5 mM EDTA; 1% Triton X-100; 1x Protease inhibitor cocktail) and two washing steps with FLAG elution buffer (20 mM Tris-HCl, pH 7.5; 150 mM NaCl; 0.2 mM EDTA; 0.1% Triton X-100; 15% glycerol), FLAG-tagged N4BP1 was eluted from M2 resins with FLAG peptide at concentration 3 μg/ml. Thirty microliters of eluate were incubated with 1 μg of either active or inactive (C360S) 6xHIS-tagged CASP8 cleavage assay buffer (50 mM HEPES, pH 7.2; 50 mM NaCl; 10 mM EDTA; 5% glycerol; 10 mM DTT) at 37**°**C for 2 h.

**GST and MBP pull-down assays**

Pull-down assays were performed as described in ^6^. Since the size of FLAG-N4BP1 (613-893) protein was identical to the size of GST diUb, GST diUb beads were additionally cleaved with 1U thrombin (*GE Healthcare*) in 1x thrombin cleavage buffer (20 mM TrisCl, pH 8.4; 150 mM NaCl; 2.5 mM CaCl_2_; 1 mM DTT) at 25**°**C for 4 h after GST PD wash.

**Pull-down of TNFR1-SC**

For isolation of TNFR1-SC complex, cells were stimulated in the presence or absence of HS-TNFα at concentration 1 μg/ml for 5 min. Then, cells were washed twice with ice-cold 1xPBS, lysed in TNFR1-SC lysis/PD buffer (20 mM Tris-HCl, pH 7.5; 150 mM NaCl, 1% Triton X-100; 10% glycerol; 2 mM NEM, 1x Protease inhibitor cocktail, 1x Phosphatase inhibitor cocktail) and incubated at 4°C for 30 min. Lysates were cleared by centrifugation (15000 rpm, 4°C, 30 min) and supernatants precleared with Superflow resin (*IBA GmbH*) at 4°C for 30 min with rotation. HS-TNFα (1 μg) was added to non-stimulated control and lysates were incubated with prewashed Strep-tactin XT resin (*IBA GmbH*) for 2 h at 4°C. Next, samples were washed 7 times with TNFR1-SC lysis/PD buffer, re-suspended in 1xLDS buffer supplemented with β-Me and denatured at 70°C for 10 min.

**Ubiquitin-binding assays**

One microgram of monoUb and synthetic diUb chains (*UbiQ Bio*) were incubated with indicated GST protein fusions bound to Glutathione Sepharose 4B resin in incubation buffer (50 mM HEPES pH 7.5; 150 mM NaCl; 1 mM EDTA; 1 mM EGTA; 1% Triton X-100; 10% glycerol and 1 mM DTT) for 2 h at 4°C. Next, samples were washed four times with incubation buffer prior to elution in 1xLDS buffer supplemented with β-Me.

**Surface plasmon resonance (SPR)**

The interactions of TF-N4BP1 (613-893) with monoUb and M1-, K48- and K63-linked diUb were analysed by SPR, using a Biacore S200 (*GE Healthcare*). Experiments were performed in HBS-P^+^ buffer (10 mM HEPES, pH 7.4, 150 mM NaCl, 0.05% Tween 20) at 25°C. TF-N4BP1 (613-893) (10 μg/ml) was covalently immobilised on a CM5 chip using the amine-coupling kit (*GE Healthcare*) according to the manufacturer’s instructions. Mono- and diUb proteins were dialyzed into HBS-P^+^ buffer prior to the experiments. Experiments were run with a concentration series of mono- and diUb at 30 μl/min with 20 s association and 30 s dissociation phases. Association and dissociation kinetics were too fast to be resolved in these experiments. Data analysis was therefore performed by analysing the plateau levels. K_D_ values were obtained from non-linear least-square fitting using a hyperbolic binding equation in GraphPad Prism 8.

**Immunoprecipitation experiments**

For HA IP, cells were washed twice in ice-cold 1x PBS, lysed in HA lysis/IP buffer (20 mM Tris-HCl, pH 7.5; 150 mM NaCl; 0.5% sodium deoxycholate; 1% NP-40; 2 mM NEM; 1 mM PMSF; 1x Protease inhibitor cocktail) and incubated with benzonase endonuclease (4°C, 30 min). Lysates were centrifuged (13000 rpm, 4°C, 15 min), followed by Sepharose CL-4B (*Sigma-Aldrich*) preclearing at 4°C for 30 min. Then, lysates were incubated with prewashed monoclonal anti-HA agarose (clone HA-7, *Sigma-Aldrich*) for 3 h at 4°C, with agitation. Samples were washed three times with denaturing buffer and twice with 1x PBS, re-suspended in 1xLDS buffer supplemented with β-Me and denatured at 70°C for 10 min.

The procedure for FLAG IP was described in *in vitro* CASP8 cleavage assay protocol. Linear Ub IP was performed as described previously^7^.

For endogenous IP cells were washed twice with ice-cold 1x PBS and lysed in endogenous IP buffer (20 mM Tris-HCl, pH 7.5; 150 mM NaCl; 1% Triton X-100; 2 mM NEM; 1x Protease inhibitor cocktail, 1x Phosphatase inhibitor cocktail) on ice for 30 min. Lysates were collected and centrifuged (13000 rpm, 4°C, 15 min), followed by incubation with the antibody recognizing desired protein and protein A/G Sepharose (*Sigma-Aldrich*) at 4°C for either 4 h or overnight. Then, samples were washed four times with an endogenous IP buffer, and eluted by incubation in a 1xLDS buffer supplemented with β-Me (70°C, 10 min).

**Dimer formation assay in cells**

HEK293T cells transfected with various wild-type and N4BP1 mutants were lysed in mild buffer containing 50 mM Tris-HCl (pH 7.5), 150 mM NaCl, 10% glycerol, 1 mM EDTA, 1% NP-40, PMSF, NEM and 1x Protease inhibitor cocktail and incubated on ice for 30 min. Lysates were centrifuged (13000 rpm, 4°C, 15 min) and either mixed with 1xLDS lacking β-Me (without 70°C for 10 min) or mixed with 1xLDS buffer supplemented with β-Me at 70°C for 10 min. Fresh lysates were immediately analysed by SDS-PAGE. For comparison, cells were also lysed in a more stringent lysis buffer (50 mM Tris-HCl, pH 7.5, 150 mM NaCl, 1% NP-40, 0.5% sodium deoxycholate, PMSF, NEM and 1x Protease inhibitor cocktail), when indicated.

**Immunoblotting**

Immunoblotting was performed as in ^6^. For linear Ub IP, immunoblotting was performed as described previously^7^.

**Preparation of IP samples for MS analysis**

For identification of cleavage sites in N4BP1, cells were transfected with plasmids encoding either FLAG-N4BP1 or N4BP1-FLAG. Twenty-four hours later, cells were washed twice in ice-cold 1x PBS, followed by lysis in denaturing buffer (20 mM Tris-HCl, pH 7.5; 150 mM NaCl; 1mM EDTA; 0.5% NP-40; 0.5% sodium deoxycholate; 0.5% SDS; 1 mM DTT; 2 mM NEM; 1x Protease inhibitor cocktail; 1x Phosphatase inhibitor cocktail). After benzonase treatment (4°C, 30 min), lysates were cleared by centrifugation (13000 rpm, 4°C, 15 min). Next, lysates were incubated with prewashed M2 resins (*Sigma-Aldrich*) for 5 h at 4°C, with agitation. Samples were washed three times with denaturing buffer and twice with distilled water and re-suspended in 1xLDS buffer supplemented with β-Me and denatured at 70°C for 10 min.

**Mass spectrometry analysis of N4BP1 cleavage**

After elution and denaturation, samples were resolved by SDS-PAGE and gel lanes were cut into 7 slices, reduced with 200 μl of 10 mM DTT, alkylated with 200 μl of 55 mM chloroacetamide and digested with trypsin (final concentration 20 μg/ml) at 750 rpm, 37**°**C, overnight. Peptides were bound to C_18_ StageTips and separated on EASY-nLC 1000 UHPLC (*Thermo Fisher Scientific*) connected to Q-Exactive HF Hybrid Quadrupole-Orbitrap (*Thermo Fisher Scientific*) mass spectrometer. For peptide separation, 15 cm and 75 μm ID PicoTip fused silica emitters (*New Objective*) were used. Emitters were self-made packed with ReproSil-Pur C18-AQ 3 μm resin (*Dr. Maisch GmbH*). Elution of the peptides from the column was performed using a linear gradient of 7-38% solvent B (80% acetonitrile in 0.1% formic acid) in 20 min with subsequent increase up to 95% solvent B within 5 min, followed by re-equilibration to 5% solvent B. Mass spectrometer was operated in positive ion mode and MS spectra were acquired with following settings: a maximal injection time of 20 ms and a 60,000/15,000 resolution at 200 m/z. Up to 15 most intense ions were selected for collision induced dissociation (CID) fragmentation. Data analysis was performed by using the MaxQuant software suite (version 1.5.3.30) and the internal search engine Andromeda and searched against the Uniprot *Homo sapiens* (released 2016) database. For the identification of cleavage sites in N4BP1, semi-specific tryptic peptides were searched for. Oxidation (M) and acetylation (protein N-terminus) were searched as variable modifications, whereas Cys carbamidomethylation (C) was set as fixed modification. Initial precursor mass tolerance was set to 4.5 ppm and MS/MS mass tolerance to 0.5 Da. Peptide and protein FDR (false discovery rate) was defined to 1%.

**Subcellular fractionation**

The cellular fractionation was performed as in ^6^.

**Luciferase assay**

The assay was performed as in ^6^. Measurements were done by using either a Wallac Victor^3^ 1420 Multilabel plate reader (*Perkin Elmer*) or Synergy H1 Hybrid Multi-mode reader (*BIOTEK*). All experiments were done at least in biological quadruplicate, where each biological replicate consisted of technical duplicate.

**Real-time PCR**

The quantitative real-time PCR was performed with SensiMix SYBR & Fluorescein kit (*Bioline*) in the iCycler iQ5 Real-Time PCR Detection System (*Bio-Rad*). GAPDH was used as an internal control. The Comparative Ct (Threshold Curve) method was used for the quantification of the amount of target, normalized to internal control. List of oligonucleotides is available in the **Supplementary table 3**. Experiment was performed in biological triplicates, where each biological replicate consisted of technical duplicates.

**Immunofluorescence**

Cells were seeded on glass coverslips and treated as indicated in the text. Cells were fixed by using 2% PFA, permeabilized with PBS containing 0.2% Triton X-100, blocked in 5% BSA in PBS solution at RT. Next, cells were incubated with appropriate antibodies. Coverslips were mounted with Mowiol containing DAPI, in order to visualize nuclei. Images were acquired with LEICA TCS SP8 confocal laser microscopy. For quantification, at least 200 cells were counted per condition.

**Crystal violet assay**

The assay was performed as in ^9^. Optical density at 570 nm (OD_570_) was measured by using Wallac Victor^3^ 1420 Multilabel plate reader (*Perkin Elmer*). Experiment was performed in biological triplicates, where each biological replicate consisted of technical quadruplicates.

**Luminescent cell viability assay**

Cell viability was determined by using CellTiter-Glo Luminescent Cell Viability Assay kit (*Promega GmbH*) according to manufacturer’s instructions. Briefly, 96-well plate was equilibrated at RT for 30 min, followed by addition of 100 μl of CellTiter-Glo Reagent provided by the manufacturer. Cell lysis was initiated by shaking a 96-well plate for 2 min. To achieve signal stabilization, 96-well plate was additionally incubated at RT for 10 min. Measurement was performed by using Synergy H1 Hybrid Multi-mode reader (*BIOTEK*). Experiments were performed in biological triplicates, where each biological replicate consisted of technical duplicates.

**NMR spectroscopy**

All NMR samples (300-500 μM) were prepared in 20 mM phosphate buffer (pH 7.0), 50 mM NaCl, 1 mM DTT, 10% D_2_O. For atom assignments, N4BP1 CUE domain was uniformly ^15^N,^13^C-labelled and assignments were completed using standard triple-resonance assignment methodology^10^. A total of 97% of the potential backbone (disregarding the proline residues) and 87% of the potential side-chain resonances were assigned (the first 3 N-terminal residues from the tag are ignored). Titration experiments involving ^15^N-labelled N4BP1 CUE domain were performed by addition of up to 5 molar equivalents of unlabeled Ub. Titration experiments involving ^15^N-labelled Ub were performed by addition of up to 5 molar equivalents of unlabeled N4BP1 CUE domain. The magnitude of chemical shift perturbations (CSPs) for each resonance was quantified according to the equation Δ𝛿 = ((𝛿^H^ _bound_-𝛿^H^_free_)^2^ + ((𝛿^N^ _bound_-𝛿^N^_free_)/a)^2^)^1/2^, where a = (𝛿^N^_max_- 𝛿^N^_min_)/ (𝛿^H^_max_- 𝛿^H^_min_). NMR experiments were performed on two types of Bruker spectrometers, an AvanceNEO 600 equipped with a 5mm ^1^H/^13^C/^15^N inverse triple resonance probe and an Avance III HD 700, equipped with a 5mm ^1^H/^13^C/^15^N triple-resonance PFG cryoprobe. All spectra were collected at 303.15 K. Data was processed using NMRPipe^11^ and analyzed using CcpNmr Analysis V2^12^.

**NMR structure determination**

An experimentally guided model of N4BP1 CUE domain was generated with NMR chemical shift data in combination with homologous structural information using the standard CS-Rosetta method^13,14^. Backbone chemical shift data (C^α^, C^β^, C’, N, H^α^ and HN) was included and a total of 20,000 models were generated. The top 10 models with the lowest energy were chosen as the final ensemble. Structural statistics were calculated using several servers including wwwPDB, MolProbity and PROSESS. Favorable Ramachandran statistics were observed, with 100% of residues in most favored (98%) regions and 0% in outlier regions (**Supplementary table S4)**.

**Molecular modelling**

The dimer of the C-terminal portion for the mouse sequence of N4BP1 was modelled with Robetta^15^ (Comparative Modelling mode). The modelled region encompasses the RNase and CUE domains including their joining linker (residues 613-893 in the UniProt sequence Q6A037). The X-ray structure of the MCPIP1 dimer (PDB ID: 5H9W) was used as template for the dimer of the RNase domains (sequence identity = 52%), while the NMR structure from this work was used for CUE domain. Sequence alignments were generated using PRALINE^16^. Multiple models were generated, which showed high variability in the relative arrangement of the CUE domains with respect to each other and to the RNase domains. This was consistent with the partially disordered and thus highly flexible nature of the linker domain (residues 776-849) as predicted by DISOPRED3^17^. Correspondingly, the estimates of model local error were generally low for the RNase domain (1.2 Å on average) and high for the linker (> 20 Å). The best model was selected to have a distance between the centres of mass of the two CUE domains compatible with a simultaneous binding to M1-linked diUb (~ 31 Å).

A model of the CUE/monoUb interface was built using HADDOCK2.4^18^ with default parameters. The structure of monoUb was taken from PDB ID: 1UBQ. CSP values were used to define the Ambiguous Interaction Restraints for the calculation. In particular, residues with CSP values greater than the average value calculated over each molecule and with a relative solvent accessible surface area (SASA) larger than 30% were set as active residues. SASA values were calculated using GetArea^19^. The solution with the best HADDOCK score (Z-score = -1.3) was also the one most consistent with the interface model emerging from the experimental data from this work and in particular with the involvement of the nonpolar interface formed between the hydrophobic patch surrounding I44 of Ub and the FP motif of the CUE domain, as well as the polar contact between K48 of Ub and D893 of N4BP1.

The final model of the N4BP1 dimer bound to the M1-linked diUb was built with MODELLER 9.15^20^. A template of the CUE/diUb complex was built by superimposing a copy of the CUE/monoUb best model from HADDOCK on each Ub molecule in the experimental structure of M1-linked diUb (PDB ID: 2W9N). The best Robetta structure (see above) was used as a template for the N4BP1 dimer. For each MODELLER run, 100 structures were generated and the one with the lowest DOPE score was selected as the final structure.

The resulting N4BP1/M1-linked diUb model was refined by energy minimisation using GROMACS 2016.3^21^. The system was solvated using a truncated octahedral box of TIP3P water molecules. A minimal distance of 12 Å was set between the protein and the walls of the box. The proteins were described with the Amber99SB*-ILDN^22^ force field. The charge of the ionisable residues was set to that of their standard protonation state at pH 7, the systems were then neutralised by adding counter-ions. Each system was minimised through 3 stages with 7000 (positional restraints on heavy atoms) + 5000 steps of steepest descent, followed by 2000 steps of conjugate gradient. The quality of the refined models was evaluated using MolProbity^23^. The refined model had a MolProbity score <=1.62 (92^nd^ percentile) and a clashscore <= 0.7 (99^th^ percentile).

**Statistical analysis**

To determine statistical significance in **Extended Figure 7D**, an unpaired, two-tailed Student’s *t* test was used. Three independent experimental replicates consisting of technical duplicates were performed. To determine statistical significance in **Extended Figure** **7B**, a two-way ANOVA test was used. Five independent experimental replicates consisting of technical duplicates were performed. To determine statistical significance in **Figures 3E, 4A, 4C** and **Extended Figure 7E**, a two-way ANOVA test, *post hoc* Sidak's multiple comparisons test were used. Three independent experimental replicates consisting of technical duplicates were performed in **Figure 3E.** Three independent experimental replicates consisting of technical triplicates were performed in **Figure 4C.** Three independent experimental replicates consisting of technical quadruplicates were performed in **Figure 4A.** To determine statistical significance in **Figure 4F**, a two-way ANOVA test, *post hoc* Tukey's multiple comparisons test were used. Three independent experimental replicates consisting of technical duplicates were performed. For all of the figures, results are shown as means and error bars defined as s.e.m. *****P* < 0.0001, ****P* < 0.001, ***P* < 0.01, **P* < 0.05 were considered significant, while *P* > 0.05 was considered nonsignificant.

**Data and material availability**

Coordinates of the structure of N4BP1-CUE and the N4BP1-CUE/Ub2 complex have been deposited in the PDB-Dev Protein Data Bank ([https://pdb-dev.wwpdb.org](https://pdb-dev.wwpdb.org/)) under accession codes PDBDEV_00000076 and PDBDEV_00000093, respectively. Chemical shift data have been deposited in the Biological Magnetic Resonance Data Bank (<https://bmrb.io>) with BMRB entry ID 50688.

The mass spectrometry proteomics data have been deposited to the ProteomeXchange Consortium via the PRIDE partner repository^8^ with the dataset identifier PXD024355.

**Extended discussion**

To confirm the dimer-dependent mechanism of Ub linkage selectivity, we compared the Ub binding specificity of N4BP1 (613-893) with or without artificially dissociated RNase domain (**Extended Data Fig. 6H**). We also tested the ability of RNase domain interface N4BP1 mutants to specifically bind to M1-linked diUb (**Extended Data Fig. 6I**). We speculated that GST-monoUb binding to individual CUE domains within the dimer would not be sterically possible, and that the low-affinity nature of monoUb interaction with N4BP1 (613-893) dimer (84.93 μM, **Table 1**) would not be detectable by GST PD. Once dimer is disrupted by stringent lysis or interface mutation, two CUE domains would separate from each other, allowing each N4BP1 (613-893) monomer to bind GST-monoUb. Indeed, we observed binding of dimerization-deficient mutants to GST-monoUb (**Extended Data Fig. 6I**), which further supports the notion that both CUE domains of the N4BP1 dimer engage in specific interaction with a single M1-linked Ub chain.

Our newly described mechanism for achieving UBD specificity therefore urges additional caution when choosing protein tags, lysing conditions and other experimental approaches during screening for and validating of novel UBDs.

### SUPPLEMENTARY REFERENCES

1.	Katoh, K., Misawa, K., Kuma, K. I. & Miyata, T. MAFFT: A novel method for rapid multiple sequence alignment based on fast Fourier transform. *Nucleic Acids Res.* (2002) doi:10.1093/nar/gkf436.

2.	Bucher, P., Karplus, K., Moeri, N. & Hofmann, K. A flexible motif search technique based on generalized profiles. *Comput. Chem.* (1996) doi:10.1016/S0097-8485(96)80003-9.

3.	Söding, J. Protein homology detection by HMM-HMM comparison. *Bioinformatics* (2005) doi:10.1093/bioinformatics/bti125.

4.	Aguileta, M. A. *et al.* The E3 ubiquitin ligase parkin is recruited to the 26 S proteasome via the proteasomal ubiquitin receptor Rpn13. *J. Biol. Chem.* **290**, 7492–7505 (2015).

5.	Pickart, C. M. & Raasi, S. Controlled synthesis of polyubiquitin chains. *Methods in Enzymology* (2005) doi:10.1016/S0076-6879(05)99002-2.

6.	Kliza, K. *et al.* Internally tagged ubiquitin: A tool to identify linear polyubiquitin-modified proteins by mass spectrometry. *Nat. Methods* **14**, 504–512 (2017).

7.	Matsumoto, M. L. *et al.* Engineering and structural characterization of a linear polyubiquitin-specific antibody. *J. Mol. Biol.* **418**, 134–144 (2012).

8.	Perez-Riverol, Y. *et al.* The PRIDE database and related tools and resources in 2019: Improving support for quantification data. *Nucleic Acids Res.* (2019) doi:10.1093/nar/gky1106.

9.	Feoktistova, M., Geserick, P. & Leverkus, M. Crystal violet assay for determining viability of cultured cells. *Cold Spring Harb. Protoc.* (2016) doi:10.1101/pdb.prot087379.

10.	Sattler, M., Schleucher, J. & Griesinger, C. Heteronuclear multidimensional NMR experiments for the structure determination of proteins in solution employing pulsed field gradients. *Progress in Nuclear Magnetic Resonance Spectroscopy* vol. 34 93–158 (1999).

11.	Delaglio, F. *et al.* NMRPipe: A multidimensional spectral processing system based on UNIX pipes. *J. Biomol. NMR* **6**, 277–293 (1995).

12.	Vranken, W. F. *et al.* The CCPN data model for NMR spectroscopy: Development of a software pipeline. *Proteins Struct. Funct. Genet.* **59**, 687–696 (2005).

13.	Shen, Y. *et al.* Consistent blind protein structure generation from NMR chemical shift data. *Proc. Natl. Acad. Sci. U. S. A.* **105**, 4685–4690 (2008).

14.	Raman, S. *et al.* NMR structure determination for larger proteins using backbone-only data. *Science (80-. ).* **327**, 1014–1018 (2010).

15.	Song, Y. *et al.* High-resolution comparative modeling with RosettaCM. *Structure* (2013) doi:10.1016/j.str.2013.08.005.

16.	Simossis, V. A. & Heringa, J. PRALINE: A multiple sequence alignment toolbox that integrates homology-extended and secondary structure information. *Nucleic Acids Res.* (2005) doi:10.1093/nar/gki390.

17.	Jones, D. T. & Cozzetto, D. DISOPRED3: Precise disordered region predictions with annotated protein-binding activity. *Bioinformatics* (2015) doi:10.1093/bioinformatics/btu744.

18.	Van Zundert, G. C. P. *et al.* The HADDOCK2.2 Web Server: User-Friendly Integrative Modeling of Biomolecular Complexes. *J. Mol. Biol.* (2016) doi:10.1016/j.jmb.2015.09.014.

19.	Fraczkiewicz, R. & Braun, W. Exact and efficient analytical calculation of the accessible surface areas and their gradients for macromolecules. *J. Comput. Chem.* (1998) doi:10.1002/(SICI)1096-987X(199802)19:3<319::AID-JCC6>3.0.CO;2-W.

20.	Šali, A. & Blundell, T. L. Comparative protein modelling by satisfaction of spatial restraints. *J. Mol. Biol.* (1993) doi:10.1006/jmbi.1993.1626.

21.	Abraham, M. J. *et al.* Gromacs: High performance molecular simulations through multi-level parallelism from laptops to supercomputers. *SoftwareX* (2015) doi:10.1016/j.softx.2015.06.001.

22.	Lindorff-Larsen, K. *et al.* Systematic validation of protein force fields against experimental data. *PLoS One* (2012) doi:10.1371/journal.pone.0032131.

23.	Chen, V. B. *et al.* MolProbity: All-atom structure validation for macromolecular crystallography. *Acta Crystallogr. Sect. D Biol. Crystallogr.* (2010) doi:10.1107/S0907444909042073.

**Extended Data Figure Legends**

**Extended Data Figure 1. N4BP1 is a novel linear ubiquitin receptor**

(**A**) GST pull-down assay with various fragments of N4BP1 transiently overexpressed in HEK293T cells with GST fusions of mono-, di- and tetraUb. GST alone was used as negative binding control. (**B**) **Top**: SeqLogo of the CUE domain consensus; amino acid frequencies have been derived from an alignment of established CUE domains. The height of each amino acid is proportional to its conservation at a respective position. **Bottom**: Alignment of N4BP1 CUE domain with CUE domain consensus, with residues invariant or conservatively replaced in at least 50% of the sequence shown on black or grey background, respectively. (**C**) Alignment of N4BP1 UBA domain with classical UBA domains. The alignment is rendered by box shade (residues invariant or conservatively replaced in at least 50% of the sequences are shown on black or grey background, respectively).

**Extended Data Figure 2. ITC analysis of the interaction of the CUE domain with different ubiquitin linkages and the RNAse domain**

Isothermal titration calorimetry was used to measure the affinity of N4BP1 (850-893) for monoUb (**A**), M1-diUb (**B**), K63-diUb (**C**), K48-diUb (**D**), monoUb (K48R) (**E**) and N4BP1 (613-773) (**F**).

**Extended Data Figure 3. NMR titrations of ^15^N-labelled N4BP1 CUE domain with different ubiquitin linkages**

(**A**) Overlay of ^1^H-^15^N-HSQC spectra of the N4BP1 (850-893) with increasing amounts of either monoUb, M1-diUb, K63-diUb or K48-diUb. Peaks are coloured according to the Ub:N4BP1 ratio of the titration. (**B**) Perturbed surface of the CUE domain upon binding to the corresponding Ub linkage. Residues with strong chemical shift perturbations are indicated. Red gradient indicates the intensity of the observed chemical shift perturbations (Δδ). ^1^H-^15^N-HSQC signals, which were completely broadened, were set to a maximum value of 0.5 ppm.

**Extended Data Figure 4. NMR titrations of ^15^N-labelled ubiquitin linkages and the CUE domain of N4BP1**

(**A**) Overlay of ^1^H-^15^N-HSQC spectra of monoUb, M1-diUb, K63-diUb and K48-diUb with increasing amounts of the N4BP1 (850-893). Peaks are coloured according to the Ub:N4BP1 ratio of the titration. (**B**) Perturbed surface of different Ub linkages upon binding to N4BP1 (850-893). The chemical shift perturbations for each residue were mapped onto the surface of monoUb (PDB ID: 1UBQ), M1-diUb (PDB ID: 2W9N), K63-diUb (PDB ID: 3H7P) and K48-diUb (PDB ID: 1ZO6). Residues with strong chemical shift perturbations are indicated. Red gradient indicates the intensity of the observed chemical shift perturbations (Δδ). ^1^H-^15^N-HSQC signals, which were completely broadened, were set to a maximum value of 0.3 ppm (prox.: proximal Ub; dist.: distal Ub). (**C**) Overlay of an expanded region from the ^1^H-^15^N-HSQC spectra of monoUb, M1-diUb and K48-diUb showing amide proton and nitrogen shifts of N25, K29 and K49 in the absence and presence of N4BP1 (850-893).

**Extended Data Figure 5. Selective binding of N4BP1 to ubiquitin chains**

**(A)** GST pull-down assay with indicated recombinant N4BP1 fragments and chemically synthesized diUb chains linked by M1, K6, K11, K27, K29, K33, K48 and K63, as well as recombinant monoUb. Schematic representation of the results is shown in (**B**). Red colour depicts the lack of binding, whereas yellow and green show weak and strong binding, respectively. (**C**) GST pull down with indicated fragments of mouse N4BP1 transiently overexpressed in HEK293T cells with GST alone, GST-monoUb and GST-diUb. After pull down, GST diUb beads were additionally incubated with thrombin, due to identical size of GST-diUb and FLAG N4BP1 (613-893).

**Extended Data Figure 6. Dimer formation of N4BP1 is required for specific recognition of M1-linked ubiquitin chains**

(**A**) Overlay of elution profiles of N4BP1 (613-774) and N4BP1 (613-893) samples loaded on an analytical Superose 12 column. Green dots correspond to elution volumes of molecular weight standards, which are plotted against the standardised elution volume Ve/Vo (**B**). The molecular weights of elution peaks 1 and 2 of N4BP1 (613-893) and peak 3 of N4BP1 (613-773) are indicated as blue dots and correspond to 121 kDa and 65 kDa for peak 1-2 which indicate dimeric higher order (tetrameric) species and 65 kDa for peak 3, indicating dimeric species. **(C-D)** Same analysis as in **(A-B)** for N4BP1 (850-893). The sample was loaded on an analytical Superdex75 column. The elution peak 4 corresponds to an apparent molecular weight of 8.2 kDa which is smaller then a the a theoretical CUE domain dimer. **(E)** Effect of lysis conditions on dimer preservation of N4BP1 (613-893). (*) indicates N4BP1 (613-893) dimer that is destroyed under denaturing conditions. **(F)** Homology model of the RNase domain of N4BP1 in cartoon and surface presentation is coloured in pale and bright orange. The dimeric structure of the RNase domain of MCPIP1 (PDB ID: 5H9W) was used as a template (C(α) trace shown in grey). Residues, which participate in dimer formation in MCPIP1 and are conserved in N4BP1 are shown as sticks. Polar interactions are indicated as dashed lines. (**G**) Schematic model of the C-terminal half of N4BP1 demonstrates that specific binding with linear Ub chains depends on dimerisation of the RNase domains. (**H**) Lysates prepared as in (**E**) were subjected to GST pull-down assay with GST fusions of mono- and diUb. **(I)** GST pull-down with wild-type and mutants of C-terminal fragment of mouse N4BP1 with GST alone, GST monoUb and GST-diUb under mild conditions. As in (**E**), GST diUb beads were additionally incubated with thrombin.

**Extended Data Figure 7. N4BP1 inhibits TNFα-stimulated NFκB signaling**

(**A**) Protein levels of N4BP1 and various components of TNFR1-SC and complex II in N4BP1^+/+^ and N4BP1^-/-^ MEFs. (**B**) Effect of N4BP1 on NFκB transcriptional activity. N4BP1^+/+^ and N4BP1^-/-^ MEFs were transiently transfected with pNFκB-Luc and pUT651 plasmids encoding luciferase and β-galactosidase, respectively. After 24 hours, cells were starved for 16 hours, followed by 6 hours stimulation with TNFα (20 ng/ml). Lysates were subjected to luciferase and β-galactosidase assays. Results are shown as means and s.e.m. (n=5). ** *P* < 0.01, * *P* < 0.05, determined by two-way ANOVA test. (**C**) Effect of N4BP1 on TNFα-induced gene expression. N4BP1^+/+^ and N4BP1^-/-^ MEFs were starved for 16 hours, followed by TNFα stimulation (20 ng/ml) for indicated time periods. mRNA levels of TNFα target genes CXCL1 (left panel) and IL-6 (right panel) were determined by real-time PCR. Results are shown as means and s.e.m. (n=3). (**D**) Quantification of p65 localization in TNFα-stimulated N4BP1^+/+^ and N4BP1^-/-^ MEFs from **Figure 3B.** Three independent experimental replicates consisting of technical duplicates were performed. At least 200 cells were quantified per condition. Results are shown as means and s.e.m. (n=3). **** *P* < 0.0001, determined by two-tailed Student’s *t*-test. (**E**) Quantification of phosphorylation (left panel) and degradation (right panel) of IκBα in N4BP1^+/+^ and N4BP1^-/-^ MEFs for **Figure 3C**. Quantification was performed with the use of ImageJ software. Results are shown as means and s.e.m. (n=3). n.s., no statistically significant difference, *P* > 0.05, * *P* < 0.05, *** *P* < 0.001, **** *P* < 0.0001, determined by two-way ANOVA, *post hoc* Sidak's multiple comparisons test. (F-**G**) Western blot analysis of cellular localization of endogenous N4BP1 in MEFs (**F**) and HEK293T cells (**G**). After cellular fractionation of N4BP1^+/+^MEFs, N4BP1^-/-^ MEFs and HEK293T cells, localization of N4BP1 was determined by using anti-N4BP1 antibody, respectively. PCNA and GAPDH were used as markers of nuclear and cytoplasmic fractions, respectively. (**H**) Effect of N4BP1 on NFκB signaling pathway upon IL-1β stimulation. After 16 hours starvation, N4BP1^+/+^ and N4BP1^-/-^ MEFs were treated with IL-1β (10 ng/ml) for indicated time periods, resolved by SDS-PAGE and analyzed by Western blot with indicated antibodies.

**Extended Data Figure 8. N4BP1 negatively regulates TNFR1 signaling through linear ubiquitin binding**

**(A)** Immunoblot showing reconstitution of N4BP1^-/-^ MEFs with empty vector, HA-N4BP1 (1-893) or HA-N4BP1 (1-893, F862G/P863A). **(B)** Determination of the effect of linear Ub binding-deficient N4BP1 on phosphorylation and degradation of IκBα. After 16 hours of serum starvation, N4BP1^-/-^ MEFs reconstituted with empty vector, HA-N4BP1 (1-893) or HA-N4BP1 (1-893, F862G/P863A) were treated with TNFα (20 ng/ml) for indicated time periods. **(C)** The effect of N4BP1 on the kinetics of linear polyUb chain assembly by LUBAC. Western blot analysis of immunoprecipitated linear Ub HMW species and total cell lysates from TNFα-treated (20 ng/ml) N4BP1^+/+^ and N4BP1^-/-^ MEFs for the indicated time periods.

**Extended Data Figure 9. N4BP1 is cleaved by CASP8**

(**A**) Proteolytic cleavage of endogenous N4BP1 upon prolonged TNFα stimulation. N4BP1^+/+^ and N4BP1^-/-^ MEFs were serum starved for 16 hours, followed by TNFα stimulation (20 ng/ml) for indicated time points. (**B**) Identification of the protease-cleaving N4BP1 upon TNFα stimulation. N4BP1^+/+^ MEFs were pre-treated with general CASP inhibitor (zVAD-fmk, 20 µM), CASP8- (z-IETD-fmk, 20 µM) or CASP3-specific (Ac-DEVD-cmk, 20 µM) inhibitors for 1 hour, followed by TNFα stimulation (20 ng/ml) for 3 hours. (**C**) Analysis of N4BP1 interaction with catalytic-dead CASP8 C360S. HEK293T cells were transiently transfected with indicated plasmids. Twenty-four hours later, immunoprecipitation of HA-CASP8 C360S was performed, followed by Western blotting with indicated antibodies. (**D**) Comparison of N4BP1 proteolytic processing in TNFα-stimulated CASP8^+/+^ and CASP8^-/-^ Jurkat cells. CASP8^+/+^ Jurkat and CASP8^-/-^ Jurkat cells were transfected with N4BP1 by electroporation. Twenty-four hours later, cells were treated with TNFα (100 ng/ml) for 3 hours. (**E**) *In vitro* cleavage of N4BP1-FLAG by recombinant CASP8. N4BP1-FLAG was immunoprecipitated under denaturing conditions and eluted with 3xFLAG peptide. Eluate was divided into 3 equal aliquots and each aliquot was incubated for 3 hours either with recombinant active CASP8, catalytic-dead CASP8 (CASP8 C360S) or kept as input. (**F**) Western blotting showing TNFα-dependent processing of FLAG-N4BP1. HEK293T cells were transiently transfected with FLAG-N4BP1. Twenty-four hours later, cells were starved for 16 hours, followed by TNFα (20 ng/ml) treatment for 6 hours. (**G**) Immunoblot representing TNFα-dependent cleavage of N4BP1-FLAG. HEK293T cells were transiently transfected with N4BP1-FLAG. Samples were prepared as in (**F**).

**Extended Data Figure 10. CASP8 cleaves N4BP1 at multiple positions**

(**A**) Schematic representation of the MS approach for identification of CASP8 cleavage sites in N4BP1. HEK293T cells transiently expressing either FLAG-N4BP1 or N4BP1-FLAG were treated with TNFα (10 ng/ml) and CHX (0.5 μg/ml) for 4 hours. Samples were resolved by SDS-PAGE and further processed for MS analysis. (**B**) MS/MS spectra showing identified N4BP1 ^297^QFSLENVPEGELLPD^311^ (top panel) and ^477^QNSSCTVDLETD^488^ (lower panel) peptides obtained upon TNFα +CHX treatment. (**C**) Western blot analysis of N4BP1-FLAG and its single, double and triple cleavage site mutants in HEK293T upon TNFα stimulation (20 ng/ml, 3 h). Twenty-four hours post transfection, HEK293T cells expressing either N4BP1-FLAG (1-893), N4BP1-FLAG (1-893, D311A), N4BP1-FLAG (1-893, D488A), N4BP1-FLAG (1-893, D484/488A) or N4BP1-FLAG (1-893, D311/484/488A) were serum starved for 16 hours, followed by TNFα stimulation (20 ng/ml) for 4 hours. (**D**) GST pull-down assay of two major N4BP1 cleavage fragments transiently overexpressed in HEK293T cells with recombinant GST fusions of mono-, di- and tetraUb. GST alone was used as negative binding control. (**E**) Reconstitution of N4BP1^-/-^ MEFs with empty vector, HA-N4BP1 (1-488), HA-N4BP1 (489-893), HA-N4BP1 (1-893) and HA-N4BP1 (1-893, D311/484/488A). (**F**) Stabilization of C-terminal cleavage fragment of N4BP1 upon inhibition of 26S proteasome. HEK293T cells were transiently transfected with N4BP1-FLAG. Twenty-four hours later, cells were starved for 16 hours, followed by TNFα (20 ng/ml) and MG132 (10 μM) treatment for 4 hours.

**Extended Data Figure 11. Comparison of different LUBID interaction modes.**

(**A**) The UBAN domain of NEMO forms a parallel homodimer. The symmetric coiled-coil generates and interface which simultaneously interacts with two M1-linked Ub chains (PDB ID: 2ZVO). **(B)** The NZF of HOIL-1L and its C-terminal α-helical extension binds to the proximal and distal Ub of linear Ub chain linkages (PDB ID: 3B08). **(C)** Structural model of N4BP1 in complex with M1-diUb. Homodimerization of N4BP1 is facilitated by the RNase domains, which enables the recognition of the proximal and distal Ub of M1-linked diUb by two CUE domains.

**Supplementary table S1.** List of plasmids.

| **Plasmid** | **Gene** | **Species** | **Source** |
| --- | --- | --- | --- |
| pUT651 | *LacZ* | E. coli | Ivan Dikic |
| CASP8, pcDNA3, C360S, HA | *CASP8* | Human | Adrian Ting |
| CASP8, pDuet1, 6His | *CASP8* | Human | This study |
| CASP8, pDuet1, C360S, 6His | *CASP8* | Human | This study |
| pSG5 Large T | *Large T antigen* | Simian virus | Addgene #9053 |
| MCPIP1 (112-132)+N4BP1 (613-774) pCold-I | *MCPIP1* | Human | This study |
|  | *N4bp1* | Mouse | This study |
| MCPIP1 (112-132)+N4BP1 (613-893) pCold-I | *MCPIP1* | Human | This study |
|  | *N4bp1* | Mouse | This study |
| N4BP1 pcDNA3.1 | *N4bp1* | Mouse | Michael Kuehn |
| N4BP1 pcDNA3.1, D311A | *N4bp1* | Mouse | This study |
| N4BP1 pcDNA3.1, D488A | *N4bp1* | Mouse | This study |
| N4BP1 pcDNA3.1, D484/488A | *N4bp1* | Mouse | This study |
| N4BP1 pcDNA3.1, D311/484/488A | *N4bp1* | Mouse | This study |
| N4BP1 pcDNA3.1, F862G, P863A | *N4bp1* | Mouse | This study |
| N4BP1 (343-893) pcDNA3.1 | *N4bp1* | Mouse | This study |
| N4BP1 pcDNA3.1, FLAG | *N4bp1* | Mouse | This study |
| N4BP1 (1-893) pFLAG-CMV1 | *N4bp1* | Mouse | This study |
| N4BP1 (1-849) pFLAG-CMV1 | *N4bp1* | Mouse | This study |
| N4BP1 (1-771) pFLAG-CMV1 | *N4bp1* | Mouse | This study |
| N4BP1 (144-893) pFLAG-CMV1 | *N4bp1* | Mouse | This study |
| N4BP1 (393-893) pFLAG-CMV1 | *N4bp1* | Mouse | This study |
| N4BP1 (393-849) pFLAG-CMV1 | *N4bp1* | Mouse | This study |
| N4BP1 (393-771) pFLAG-CMV1 | *N4bp1* | Mouse | This study |
| N4BP1 (1-392) pFLAG-CMV1 | *N4bp1* | Mouse | This study |
| N4BP1 (1-143) pFLAG-CMV1 | *N4bp1* | Mouse | This study |
| N4BP1 (144-392) pFLAG-CMV1 | *N4bp1* | Mouse | This study |
| N4BP1 (1-488), NpFLAG-CMV1 | *N4bp1* | Mouse | This study |
| N4BP1 (489-893), NpFLAG-CMV1 | *N4bp1* | Mouse | This study |
| N4BP1 (613-893), NpFLAG-CMV1 | *N4bp1* | Mouse | This study |
| N4BP1 (613-893), NpFLAG-CMV1, R691A | *N4bp1* | Mouse | This study |
| N4BP1 (613-893), NpFLAG-CMV1, R697A | *N4bp1* | Mouse | This study |
| N4BP1 (613-893), NpFLAG-CMV1, K712A | *N4bp1* | Mouse | This study |
| N4BP1 (613-893), NpFLAG-CMV1, Q743A | *N4bp1* | Mouse | This study |
| N4BP1 (613-893), NpFLAG-CMV1, D755R | *N4bp1* | Mouse | This study |
| N4BP1 (613-893), NpFLAG-CMV1, Δ775-849 | *N4bp1* | Mouse | This study |
| N4BP1 (613-849), NpFLAG-CMV1 | *N4bp1* | Mouse | This study |
| N4BP1 (613-774), NpFLAG-CMV1 | *N4bp1* | Mouse | This study |
| N4BP1 (312-392) pEGFP-C1 | *N4bp1* | Mouse | This study |
| N4BP1 (312-342) pEGFP-C1 | *N4bp1* | Mouse | This study |
| N4BP1 (703-896) pEGFP-C1 | *N4BP1* | Human | This study |
| N4BP1 (850-893) pEGFP-C1 | *N4bp1* | Mouse | This study |
| N4BP1 (850-893) pEGFP-C1, F862G, P863A | *N4bp1* | Mouse | This study |
| N4BP1 (850-893) pEGFP-C1, D893A | *N4bp1* | Mouse | This study |
| N4BP1 (1-893), pcDNA3, HA | *N4bp1* | Mouse | This study |
| N4BP1 (1-893), pBabe-puro, HA | *N4bp1* | Mouse | This study |
| N4BP1 (1-893), D311A, D484A, D488A,  pBabe-puro, HA | *N4bp1* | Mouse | This study |
| N4BP1 (1-488), pBabe-zeo, HA | *N4bp1* | Mouse | This study |
| N4BP1 (489-893), pBabe-puro, HA | *N4bp1* | Mouse | This study |
| N4BP1 (850-893) pGEX-4T1 | *N4bp1* | Mouse | This study |
| N4BP1 (706-893) pGEX-4T1 | *N4bp1* | Mouse | This study |
| N4BP1 (343-893) pGEX-4T1 | *N4bp1* | Mouse | This study |
| N4BP1 (613-774) pCold-TF | *N4bp1* | Mouse | This study |
| N4BP1 (613-893) pCold-TF | *N4bp1* | Mouse | This study |
| N4BP1 (850-893), pET47 | *N4bp1* | Mouse | This study |
| hTNF (77-233)-pRSET-His-PP, STREP | *TNFα* | Human | This study |
| Ub, pGEX-4T1 | *UBB* | Human | This study |
| Ub, pGEX-6P1 | *UBB* | Human | This study |
| Ub, pGEX-6P1, K48A | *UBB* | Human | This study |
| Ub, pGEX-6P1, K48R | *UBB* | Human | This study |
| diUb, pET47 | *UBB* | Human | This study |
| diUb-pGEX-4T1 | *UBB* | Human | This study |
| diUb-pGEX-6P1 | *UBB* | Human | This study |
| tetraUb-pGEX-4T2 | *UBB* | Human | Errol Friedberg, Caixia Guo |
| hexaUb (GV, deltaGG) YTH9 | *UBB* | Human | This study |
| UBE2R1, pET15b, 6His-SUMO1 | *UBE2R1* | Yeast | MRC Protein Phosphorylation and Ubiquitylation Unit, DU No. 51247 |
| UBE2D3, pGEX-6P1 | *UBE2D3* | Human | This study |
| UBE2N, pET15b, 6His 3C, E60C | *UBC13* | Human | MRC Protein Phosphorylation and Ubiquitylation Unit, DU No. 24653 |
| UBA1, pET28 | *UBA1* | Mouse | This study |
| UBE2V1, pET15b, 6His 3C | *UBE2V1* | Human | MRC Protein Phosphorylation and Ubiquitylation Unit, DU No. 20496 |

**Supplementary table S2.** List of antibodies.

| **Protein/tag** | **Antibody** | **Source** | **Application** |
| --- | --- | --- | --- |
| CASP3, cleaved | 5A1E (9664) | Cell Signaling | WB |
| CASP8 | D35G2 (4790) | Cell Signaling | WB |
| FADD | sc-6036 | Santa Cruz Biotechnology | IP, WB |
| GAPDH | 14C10 (2118) | Cell Signaling | WB |
| IκBα | 112B2 (9247) | Cell Signaling | WB |
| N4BP1 (mouse-specific) | ab133610 | Abcam | IP, WB |
| N4BP1 (human-specific) | ab169329 | Abcam | WB |
| p65 | C-20 (sc-372) | Santa Cruz Biotechnology | IF |
| PARP | 9542 | Cell Signaling | WB |
| Phospho-p38 | 9216 | Cell Signaling | WB |
| Phospho-IκBα | 5A5 (9246) | Cell Signaling | WB |
| Phospho-JNK | 9255 | Cell Signaling | WB |
| PCNA | PC10 (sc-56) | Santa Cruz Biotechnology | WB |
| RIP1 | 610459 | BD Transduction Laboratories | IP, WB |
| TRADD | sc-7868 | Santa Cruz Biotechnology | WB |
| TRAF2 | 4724 (C192) | Cell Signaling | WB |
| Ubiquitin (linear) | 1F11/3F5/Y102L | Genentech | IP, WB |
| Ubiquitin | 3933 | Cell Signaling | WB |
| Ubiquitin | P4D1 (sc-8017) | Santa Cruz Biotechnology | WB |
| FLAG | M2 | Sigma-Aldrich | IP, WB |
| GFP | sc-9996 | Santa Cruz Biotechnology | WB |
| HA | HA.11 | Covance | WB |
| STREP | 34850 | Qiagen | WB |
|  | goat anti-mouse IgG HRP conjugate | Bio-Rad | 2^nd^ Ab |
|  | goat anti-rabbit HRP conjugate | DAKO | 2^nd^ Ab |
|  | donkey anti-goat HRP conjugate | Santa Cruz Biotechnology | 2^nd^ Ab |
|  | goat anti-human HRP conjugate | Jackson ImmunoResearch | 2^nd^ Ab |
|  | donkey anti-rabbit Cy5 conjugate | Jackson ImmunoResearch | IF |

WB=Western blot, IP=immunoprecipitation, 2^nd^ Ab=secondary antibody

**Supplementary table S3.** List of oligonucleotides used for RT PCR.

| **Oligonucleotide** | **Sequence (5´-3´)** | **Company** |
| --- | --- | --- |
| *Gapdh,* Sense | ACCACAGTCCATGCCATCAC | *Eurofins Genomics* |
| *Gapdh,* Antisense | CACCACCCTGTTGCTGTAGCC | *Eurofins Genomics* |
| *Cxcl1,* Sense | GCCTATCGCCAATGAGCTG | *Eurofins Genomics* |
| *Cxcl1,* Antisense | TGGGGACACCTTTTAGCATC | *Eurofins Genomics* |
| *Il-6, Sense* | CCGGAGAGGAGACTTCACAG | *Eurofins Genomics* |
| *Il-6, Antisense* | GGAAATTGGGGTAGGAAGGA | *Eurofins Genomics* |

**Supplementary table S4.** Summary of the structure statistics for the N4BP1 CUE CS-Rosetta model.

| Experimental restraints input for CS-Rosetta | |
| --- | --- |
| ^13^C^α^ shifts | 44 |
| ^13^C^β^ shifts | 44 |
| ^13^C’ shifts | 43 |
| ^15^N shifts | 40 |
| ^1^H^N^ shifts | 40 |
| ^1^H^α^ shifts | 44 |
| Total restraints | 255 |
| Average pairwise RMSD* (Å) | |
| C^α^ | 0.49 |
| Backbone atoms | 0.48 |
| Heavy atoms | 0.80 |
| All atoms | 1.01 |
| Structure quality** | |
| Ramachandran favoured regions (%) | 100 |
| Ramachandran Outliers (%) | 0 |
| Ramachandran distribution Z-score | |
| Whole | -1.43 ± 0.90 |
| Helix | -0.69 ±0.60 |
| Sheet | None |
| Loop | -1.64 ± 1.39 |
| Favored rotamers (%) | 100 |
| Poor rotamers (%) | 0 |
| C^β^ deviations >0.25 Å | 0 |
| Bad bonds (%) | 0 |
| Bad angles (%) | 0 |
| All-atom clashscore*** | 1.4 |

* Pairwise RMSD was calculated using the structure ensemble containing 10 best refined models.

**Quality data for model 5 from the Structure ensemble. Model 5 is the overall representative, medoid model (most similar to other models in the ensemble).

***Clashscore is the number of serious steric overlaps (>0.4 Å) per 1000 atoms.
